# Insulin receptor turnover in fasting is dependent on β-dystroglycan deglycosylation

**DOI:** 10.1101/2022.06.24.497215

**Authors:** Sunu Joseph, Sewar Zbidat, Alexandra Volodin, Dharanibalan Kasiviswanathan, Adina I. Fried, Andrea Armani, Jennifer E. Gilda, Shenhav Cohen

## Abstract

Fasting exerts various physiological effects, most notably, reduced signaling through the insulin receptor. We showed that insulin receptor activity requires association with Dystrophin Glycoprotein Complex (DGC). Here, we demonstrate that insulin receptor turnover by lysosomes during fasting is dependent on deglycosylation of the principal DGC component, β-dystroglycan. We show that the lysosomal enzymes HexA and Man2b1, which specifically remove N-linked glycans, mediate β-dystroglycan deglycosylation and consequently insulin receptor-DGC loss. Surprisingly, the lysosomal enzyme NAGLU, which cannot process N-linked glycosylation, also facilitated β-dystroglycan deglycosylation and insulin receptor loss. NAGLU enhances the activity of the transcriptional complex PPAR-γ/RXR-α, which in turn promotes Man2b1 and HexA induction and the resulting β-dystroglycan deglycosylation. Accordingly, downregulation of HexA, Man2b1, NAGLU or RXR-α during fasting blocked β-dystroglycan deglycosylation, and caused accumulation of insulin receptor-DGC assemblies on the membrane. Thus, NAGLU mediates physiological adaptation to fasting by promoting indirectly β-dystroglycan deglycosylation.

## INTRODUCTION

The insulin/insulin-like growth factor 1 signaling is a nutrient-sensing pathway that plays a central role in maintenance of cellular homeostasis in all cells. By binding to its tyrosine kinase cell surface receptor, insulin simulates physiological decisions to grow, proliferate, and increase energy stores. In times of scarcity, such as during fasting, when the body needs immediate supply of glucose to nurture the brain, the conversion of amino acids to glucose by hepatic gluconeogenesis is crucial for survival. Deceased circulating insulin is required for survival because the resulting fall in insulin signaling in skeletal muscle activates protein degradation via the proteasome and autophagy, elevating the levels of circulating amino acids and their conversion to glucose by the liver. Here, we uncovered a mechanism that mediates this physiological response to fasting, which involves the rapid turnover of the insulin receptor through effects on the principle component of Dystrophin Glycoprotein Complex (DGC), β-dystroglycan.

When blood glucose levels rise, insulin is secreted from pancreatic β cells and binds its receptor to coordinate nutrient uptake and cellular energy homeostasis. The insulin receptor is composed of two subunits, α and β, which are encoded by a single *INSR* gene, and are generated by posttranslational cleavage of the linear gene product ^1^. Binding of insulin to the α subunit facilitates association with the β subunit to form a functional insulin receptor that harbors tyrosine kinase activity ^1^. As a result, the heterodimeric receptor undergoes autophosphorylation and activates PI3K-AKT signaling, which in most cells controls proliferation, and in non-dividing muscle cells stimulates protein synthesis and blocks proteolysis ^2^. In muscle, activation of insulin signaling promotes growth by suppressing the transcription factors Forkhead box protein O (FOXO), which control the expression of atrophy-related genes (named ‘atrogenes’) ^3,4^. However, during fasting, when blood insulin levels fall, the resulting suppression of insulin signaling stimulates FOXO-mediated proteolysis and muscle atrophy ^5^, and the amino acids produced are converted to glucose in the liver to nurture the brain. In addition to the fall in circulating insulin, adaptation to physiological demand during fasting may involve a reduction in insulin receptor levels on the cell surface ^6^.

Recycling of cell surface receptors is not only a mechanism to terminate signal transmission, but also to promote cargo shuttling from the plasma membrane to the endosomal compartments. Internalization of insulin receptor upon insulin binding has been proposed to regulate transmission of growth signals from the cell surface ^7^. Such internalization may be enhanced to efficiently terminate insulin signaling as an adaptive physiological response to fasting and as a pathological sequel of insulin resistance in type-2 diabetes. Accordingly, our exciting recent discoveries regarding the dynamics of insulin receptor recycling concerns increased internalization and loss during fasting or diabetes, concomitantly with internalization and degradation of the DGC ^6^. Performing biochemical analysis of mouse skeletal muscles along with high-resolution microscopy we demonstrated that insulin receptors physically and functionally interact with DGC on the plasma membrane via the desmosomal component plakoglobin ^6^. Conversely, reduced insulin signaling, as occurs in fasting or catabolic diseases (e.g. type-2 diabetes), results in loss of these protein assemblies via autophagy. Under these catabolic conditions, the autophagy marker LC3 rises and competes with plakoglobin for binding to β-dystroglycan, and through association with β-dystroglycan’s LC3-interacting region (LIR) motifs ^8^ directs the recycling vesicles containing plakoglobin-DGC-insulin receptor assemblies to the lysosome ^6^. The present studies surprisingly demonstrate that such loss of insulin receptors during fasting is driven by the deglycosylation of β-dystroglycan.

The DGC is essential for muscle architecture and function by linking the desmin cytoskeleton to the extracellular matrix (ECM) ^9,10^, and its dissociation causes dystrophies in humans including Duchenne-Muscular Dystrophy (DMD) ^11^ and Becker-Muscular Dystrophy ^12^, and largely contributes to the atrophy seen in the elderly ^13^. This multi-subunit complex is composed of dystrophin, sarcoglycans, sarcospan, dystrobrevins, syntrophin and α- and β-dystroglycan ^14,15^, and β-dystroglycan serves as the core subunit spanning the muscle membrane and linking the ECM via α-dystroglycan to dystrophin and the intracellular cytoskeleton ^16^. Mounting evidence indicate that glycosylation of α-dystroglycan is important for association with ECM proteins ^17–21^. However, the less extensively studied N-linked glycosylation of β-dystroglycan appears essential for DGC integrity ^14,22^, and perturbation of this glycosylation causes myopathies in humans ^23^ and correlates with DGC dissociation in catabolic states such as fasting or diabetes ^6^. We demonstrate here that β-dystroglycan deglycosylation in fasting is mediated by the lysosomal enzymes Mannosidase Alpha Class 2B Member-1 (Man2b1) and Hexosaminidase A (HexA), and that this deglycosylation facilitates insulin receptor internalization and trafficking to lysosomes. Man2b1 and HexA are induced by the transcriptional complex Peroxisome Proliferator-Activated Receptor-γ (PPAR-γ)/Retinoid X Receptor-α (RXR-α), whose activation requires the lysosomal enzyme alpha-N-acetylglucosaminidase (NAGLU). Thus, NAGLU functions as a novel mediator of physiological adaptation to fasting by indirectly promoting DGC-insulin receptor loss, which is critical for activation of proteolysis in muscle and consequently glucose production in the liver to nurture the brain.

## RESULTS

### In fasting, β-dystroglycan-insulin receptor assemblies accumulate in lysosomes

We recently showed that plakoglobin, insulin receptor and β-dystroglycan interact in normal muscle, and during fasting the membrane content of these co-assemblies is reduced ^6^. Consistently, plakoglobin immunoprecipitation from membrane extracts of mouse muscles indicated that while plakoglobin-insulin receptor association remained intact, the association with glycosylated β-dystroglycan was markedly reduced in fasting compared with fed control (Fig. 1a) ^6^. Because colchicine injection of mice to inhibit autophagy flux attenuated this response to fasting ^6^, we hypothesized that during fasting these protein co-assemblies are shunted to lysosomes, where they are probably degraded. To test this idea, we isolated by differential centrifugation a crude lysosomal extract from whole Tibialis Anterior (TA) muscles from fed (53.6mg TA weight) and fasted (2 d) (30.4mg TA weight) mice, and subjected the obtained samples to flotation in discontinuous low-osmolarity Nycodenz gradient (Fig. 1b). Density perturbation had been shown to provide high yield of lysosomes and to improve the resolution of the commonly used Percoll gradients ^24,25^. Crude lysosomal extracts were adjusted to high density (in 45% Nycodenz solution) and layered under the 19%-30% discontinuous density gradient, and following high-speed centrifugation organelles were separated based on their density (Fig. 1b). Bottom-collection of banded material and analysis by immunoblotting using antibodies against the lysosomal marker LAMP1 and the lysosomal enzyme cathepsin D indicated a successful fractionation of lysosomes of low (LD) and high (HD) densities (Fig. 1c). Heterogeneity of lysosomes with regard to density and enzyme content has been reported before ^26^. In agreement with stimulated autophagy and increased number of lysosomes in atrophying muscles during fasting ^27,28^, a larger amount of lysosomes (indicated by the stronger signal for lysosomal markers) was recovered in various densities across the Nycodenz gradient at 2 d of fasting compared with fed control (Fig. 1c), even though the atrophying muscle used for analysis was much smaller than the one from fed mice (30.4mg and 53.6mg TA weights from fasted and fed mice, respectively). In addition, other autophagic and endosomal organelles were enriched in the atrophying muscle including EEA1-labeled early and Rab7-labeled late endosomes, as well as LC3-II, a protein marker of autophagosomes (Fig. 1c). The high-density lysosomal particles probably emanated from low density lysosomes that progressively became denser by the build-up of biomaterial destined for degradation. In fact, the fractions containing lysosomes and autophagosomes in atrophying muscles also contained increased amounts of insulin receptor, β-dystroglycan, and plakoglobin, while the levels of these proteins on the muscle membrane were reduced (see below) ^6^, strongly suggesting that these proteins were shunted to lysosomes (Fig. 1c). In fasting, unmodified β-dystroglycan preferentially accumulated in lysosomes suggesting that β-dystroglycan deglycosylation was initiated on the muscle membrane and preceded trafficking to lysosomes (Fig. 1c graph, and see below). It is noteworthy that these preparations contained small traces of ER and Golgi as shown by antibodies against specific markers for these organelles, Calnexin and Golgin 97, respectively (Fig. 1c). Also, using an antibody against the mitochondrial marker ATP synthase-β we discovered that the four highest density fractions of the crude lysosomal extracts also contained traces of mitochondria (Fig. 1c) as previously described ^26,29,30^, most likely because mitochondria has membrane-contacts sites with lysosomes ^31,32^. These lysosome-mitochondria contacts mediate transport of metabolites such as phospholipids, sugars and amino acids generated in lysosomes for energy production in mitochondria ^33^.

**Figure 1.**
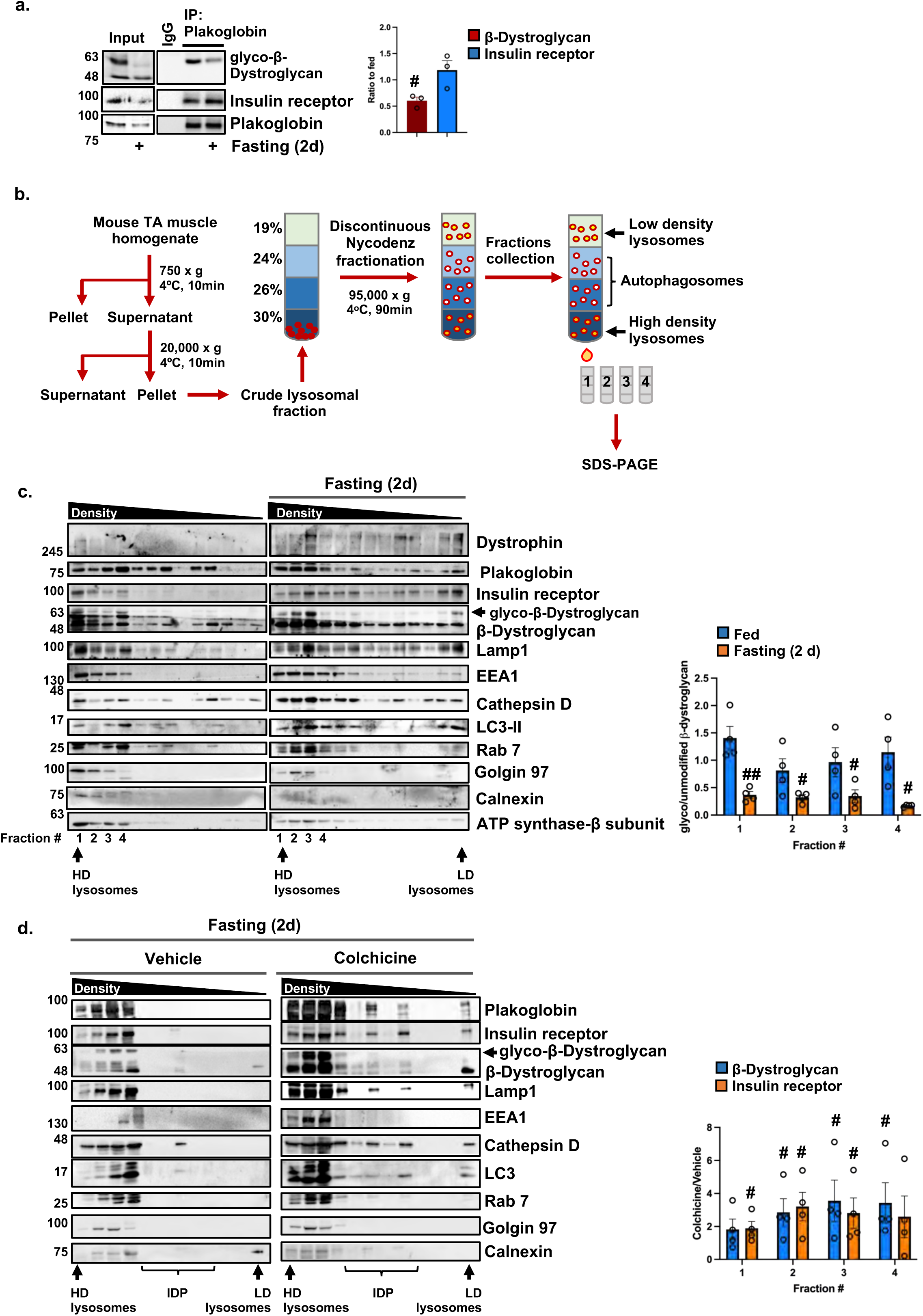
During fasting, β-dystroglycan-insulin receptor assemblies accumulate in lysosomes. (a) Plakoglobin immunoprecipitation from whole muscle membrane extracts from fed and fasted (2 d) mice, and analysis by SDS-PAGE and immunoblotting. The data are representative of three independent experiments. Graph: data were normalized to plakoglobin and plotted as the mean fold change relative to fed control ± SEM. n=3 mice per condition. #, p<0.05 *vs.* fed by one-tailed unpaired Student’s *t* test. (b) Schematic for preparation of crude lysosomal extract and floatation in Nycodenz gradient. (c) During fasting, β-dystroglycan-insulin receptor-plakoglobin assemblies accumulate in lysosomes. Crude lysosomal extracts from muscles of fed (TA weight 53.6mg) and fasted (TA weight 30.4mg) mice were subjected to flotation in a Nycodenz gradient, and fractions were analyzed by SDS-PAGE and immunoblotting. Low density (LD); High density (HD). Graph depicts densitometric measurements of β-Dystroglycan blots as glycosylated/unmodified ratio, from four independent experiments. Data are represented as mean ± SEM. n=4 mice per condition. #, p<0.05; ##, p<0.005 each fraction in fasting *vs.* fed by one-tailed unpaired Student’s *t* test. (d) Crude lysosomal extracts from muscles of fasted mice treated with colchicine or vehicle were subjected to flotation in Nycodenz gradient and analyzed by immunoblotting. Intermediate density particles (IDP). Colchicine (0.4mg/kg body weight) was injected at 24 and 36 hours after food deprivation, and mice were sacrificed at 48 hours of fasting. Graph depicts densitometric measurements of β-Dystroglycan and insulin receptor as colchicine/vehicle ratio, from four independent experiments. Data are represented as mean ± SEM. n=4 mice per condition. #, p<0.05 each fraction in colchicine *vs.* vehicle by one-tailed unpaired Student’s *t* test.

To determine if the density heterogeneity among lysosomes in fact represents different stages of one common catabolic route, we inhibited autophagy flux by injecting mice with colchicine (0.4mg/kg body weight) ^6,34,35^ every 12 hours, starting at 24 hours after food deprivation to allow trafficking of plasma membrane proteins to the lysosome during the first 24 hours of fasting. At 2 d of fasting, the mice were sacrificed and muscles analyzed by fractionation of crude lysosomal extracts on Nycodenz gradient (Fig. 1d). To make any small differences more evident, 50% of crude lysosomal extracts from muscles of mice injected with colchicine or vehicle were loaded on the Nycodenz gradient; as a result, generally less proteins were recovered in lysosomes in fasting (vehicle) compared to the fasting sample presented in Fig. 1c. Similar to our observations in Fig. 1c, lysosomes were mainly enriched in the high density fractions at 2 d of fasting (Fig. 1d), while 24 hours of fasting followed by colchicine injection led to recovery of lysosomes as intermediate density particles (IDP)(Fig. 1d). The low, intermediate, and high-density lysosomes seem to participate in the same catabolic process because plakoglobin, insulin receptor, and β-dystroglycan, which sedimented to high density Nycodenz fractions at 2 d of fasting (Figs. 1c-d), appeared also in intermediate density fractions on colchicine treatment (Fig. 1d). These proteins were degraded during fasting, when their levels on the plasma membrane were reduced ^6^, because colchicine injection of mice led to accumulation of plakoglobin, insulin receptor, and β-dystroglycan mostly in autophagosomes (detected with LC3 antibodies), which sedimented to the four highest density fractions of the crude lysosomal extracts (Fig. 1d). HD lysosomes and autophagosomes appear to have similar densities because complete separation of these two types of vesicles is known to be a technical challenge. Moreover, in the Nycodenz fractions containing lysosomes, β-dystroglycan appeared mostly in its deglycosylated form (Fig. 1c), and mechanisms that promote β-dystroglycan deglycosylation should also reduce DGC integrity and insulin receptor stability ^6,14^.

### Man2b1 and HexA mediate β-dystroglycan deglycosylation and insulin receptor loss

To identify the enzymes that are involved in β-dystroglycan deglycosylation and determine the role of β-dystroglycan deglycosylation in promoting insulin receptor loss, it was important to initially confirm that β-dystroglycan is N-linked glycosylated ^16,36^ in skeletal muscle. Therefore, we established an *in vitro* deglycosylation assay by incubating skeletal muscle homogenates with recombinant Peptide:N-glycosidase-F (PNGase-F), which specifically removes N-linked (β-GlcNAc) glycans on glycoproteins. Analysis of the enzymatic reactions by immunoblotting indicated that in the presence of PNGase-F, the high molecular weight species of β-dystroglycan, corresponding to the glycosylated protein, disappeared and β-dystroglycan accumulated in its unmodified form (Fig. 2a). These findings are consistent with prior reports showing N-linked glycosylation of β-dystroglycan ^22^. Thus, in normal muscle β-dystroglycan is N-linked glycosylated.

**Figure 2.**
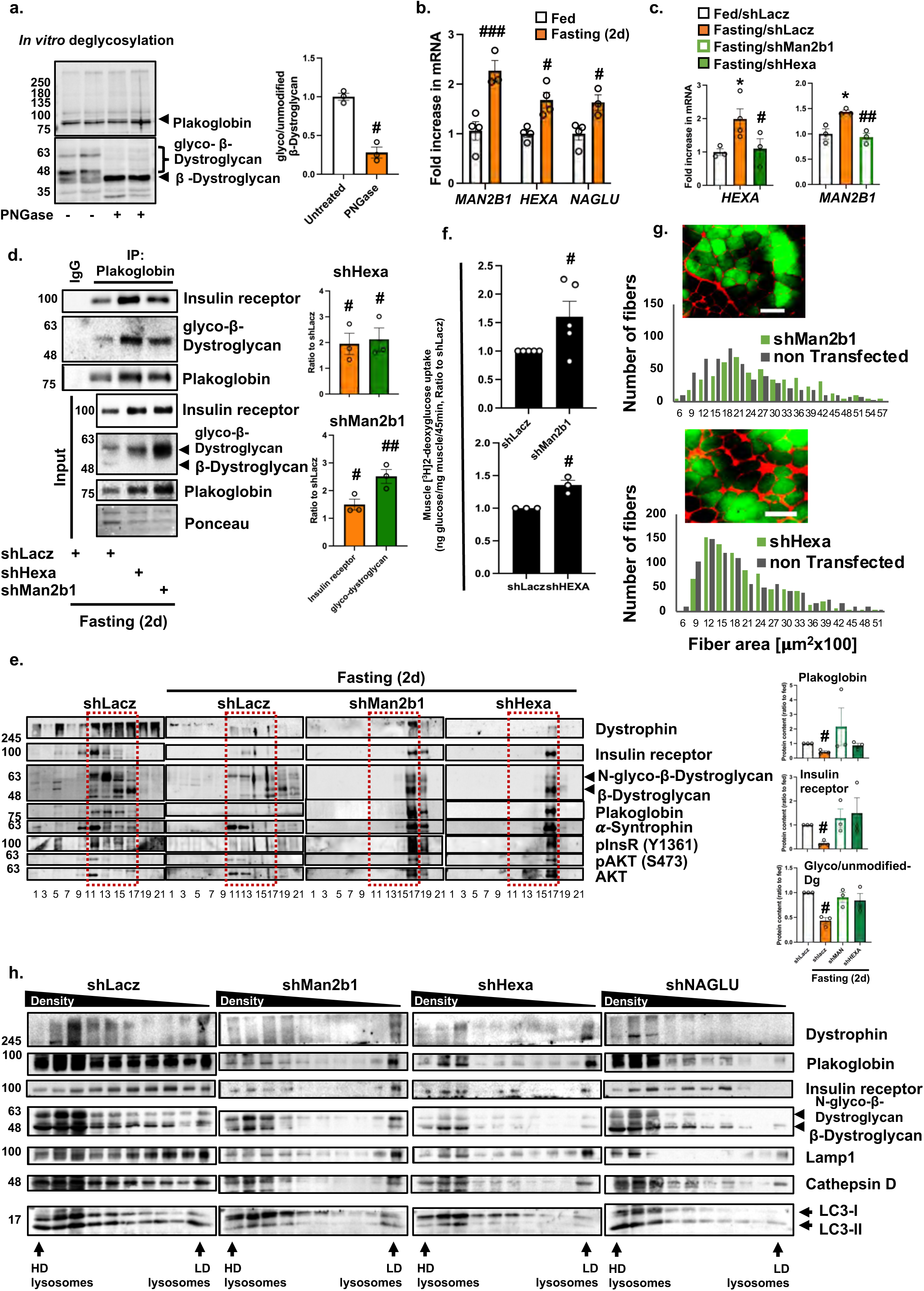
Man2b1 and HexA mediate β-dystroglycan deglycosylation and insulin receptor loss. (a) In skeletal muscle, β-dystroglycan is N-linked glycosylated. Whole muscle extracts (25 μg) were incubated with PNGase F *in vitro* and samples were analyzed by immunoblotting. The data are representative of three independent experiments. Graph depicts mean ratio to untreated ± SEM. n=3 mice. #, p<0.05 *vs.* untreated by one-tailed paired Student’s *t* test. (b) RT-PCR of mRNA preparations from normal or atrophying muscles using primers for *MAN2B1*, *HEXA*, or *NAGLU*. Data are plotted as the mean fold change relative to control ± SEM. n=3-4 mice per condition. #, p<0.05; ###, p<0.0005 *vs.* fed by two-way ANOVA. (c) RT-PCR of mRNA preparations from muscles expressing shMan2b1, shHexA, or shLacz from fed or fasted mice using specific primers for *MAN2B1* and *HEXA*. Data are plotted as the mean fold change relative to control ± SEM. n=3-4 mice per condition. *, p<0.05 *vs.* shLacz in fed and #, p<0.05; ##, p<0.005 *vs.* shLacz in fasting by ANOVA. (d-e) During fasting, downregulation of *MAN2B1* or *HEXA* promotes accumulation of intact glycosylated β-dystroglycan-insulin receptor-plakoglobin assemblies on the muscle membrane. Membrane extracts from whole TA muscles expressing shMan2b1, shHexA, or shLacz were subjected to plakoglobin immunoprecipitation (d) or glycerol gradient fractionation (e), and analyzed by SDS-PAGE and immunoblotting. Ponceau S staining is shown as a loading control for input blots. The data are representative of three independent experiments. (d) Graphs depict each protein relative to plakoglobin as a ratio to shLacz. Data are mean ± SEM. n=3 mice. #, p<0.05 and ##, p<0.005 *vs.* shLacz by one-tailed unpaired Student’s *t* test. (e) densitometric measurement of fraction #17 in all gradients. Data are mean ± SEM. n=3 mice per condition. #, p<0.05 *vs.* shLacz in fed by one-way ANOVA. (f) In vivo [^3^H]-2-deoxyglucose uptake by muscles expressing shLacz, compared to the contralateral limbs expressing shHexa or shMan2b1, is plotted as ratio to shLacz. Data represent ng glucose/mg muscle/45 min as mean ± SEM. n = 3-5 mice. # P < 0.05 *vs.* shLacz by one-tailed paired Student’s *t* test. The data are representative of two independent experiments. (g) Measurements of cross-sectional areas of 527 fibers expressing shMan2b1 or 821 fibers expressing shHexA (green bars) *vs.* the same number of adjacent non-transfected fibers (dark bars). n=4 mice. Dystrophin staining is in red. Bar, 50μm. (h) During fasting, downregulation of *MAN2B1*, *HEXA* or *NAGLU* prevents accumulation of β-dystroglycan-insulin receptor-plakoglobin in lysosomes. Crude lysosomal extracts from muscles expressing shMan2b1, shHexA, or shNAGLU from fasted mice were subjected to flotation in Nycodenz gradient and analyzed by SDS-PAGE and immunoblotting. The data are representative of four independent experiments.

Because β-dystroglycan deglycosylation reduces DGC integrity ^14^, which in fasting it is coupled to insulin receptor loss ^6^, it stands to reason that these processes would be causally linked. Using ectopically expressed 6His-tagged plakoglobin and mass spectrometry ^6^, we identified three lysosomal glycoside hydrolases (2 unique peptides for each) that bound plakoglobin-insulin receptor-β-dystroglycan co-assemblies in muscle homogenates, including Man2b1 and HexA, which specifically catalyze the release of N-linked glycans during glycoprotein turnover, plus a α-N-acetylglucosaminidase (α-GlcNAc-specific hydrolase) called NAGLU. These two types of glycosylations are highly attuned to nutrient availability because they rely on glucose metabolism ^37^. Accordingly, on food deprivation, when circulating glucose is low and β-dystroglycan on the muscle membrane is deglycosylated (Fig. 1a and ^6^), Man2b1, HexA, and NAGLU were induced in atrophying muscles (Fig. 2b).

Because Man2b1 and HexA specifically remove N-linked glycosylation, we determined whether they contribute to β-dystroglycan deglycosylation during fasting. To clarify their roles, we suppressed Man2b1 and HexA expression by electroporation into mouse TA of specific shRNA plasmids designed against the mouse genes (shMan2b1 and shHexA), which completely prevented the induction of Man2b1 and HexA in atrophying muscles (Figs. 2c and S1a-b). The downregulation of Man2b1 and HexA during fasting resulted in a marked increase on the muscle membrane of glycosylated β-dystroglycan compared to muscles expressing shLacz control (Fig. 2d, input). These findings were corroborated by immunofluorescent staining of transfected muscles, where β-dystroglycan accumulated on the muscle membrane in fibers expressing shMan2b1 or shHexA (also express GFP)(Fig. S1c). Moreover, this increase in the amount of glycosylated β-dystroglycan was completely reversed by the ectopic co-expression of Myc-tagged full-length Man2b1 or HexA in atrophying muscles (Fig. S1a-b), indicating that the deglycosylation of β-dystroglycan during fasting is specifically mediated by these enzymes.

In addition, the accumulation of glycosylated β-dystroglycan in muscles expressing shMan2b1 or shHexA was accompanied by an accumulation of insulin receptor and plakoglobin, which were associated with each other on the muscle membrane in one intact complex because they co-precipitated with plakoglobin from membrane extracts (Fig. 2d). Consistently, glycerol gradient fractionation of membrane extracts indicated increased amounts of glycosylated β-dystroglycan-insulin receptor-plakoglobin co-assemblies in muscles where Man2b1 or HexA were downregulated relative to shLacz control (Fig. 2e). In these muscles, glycosylated β-dystroglycan, insulin receptor, and plakoglobin sedimented to the same glycerol gradient fractions (Fig. 2e, marked by a red rectangle) as an intact co-assembly (Fig. 2d). This stabilization of insulin receptor-DGC on the membrane promoted insulin receptor activity and PI3K-AKT signaling because the levels of phosphorylated insulin receptor and AKT were increased during fasting compared to shLacz expressing controls (Fig. 2e), which was sufficient to enhance glucose uptake (Fig. 2f). It is noteworthy that in these muscles, the membrane content of the protein co-assembly was lower than that in fed mouse muscles (Fig. 2e), probably because the atrophying muscles were about 70% transfected with shMan2b1 or shHexA.

Our previous studies suggested that β-dystroglycan-insulin receptor-plakoglobin association is important for maintenance of muscle fiber size ^6^. Consistently, the decrease in Man2b1 and HexA levels by specific shRNAs was sufficient to attenuate fiber atrophy during fasting because the cross-sectional area of fibers expressing shManb1 or shHexA was significantly larger than that of non-transfected fibers (Fig. 2g and Table 1). To quantify statistically the effects of shManb1 and shHexA on fiber size, we adopted the Vargha-Delaney A-statistic test, which is defined as “the probability of x ≥ y, where x, y are random variables sampled from distributions X and Y” ^38^ (Table 1). As indicated in our recent paper ^38^, the A-statistic is a direct measure of effect on size, and it takes on values between 0 and 1, with 0 ≤ A < 0.5 indicating X is stochastically less than Y. Our data show that the A-statistic values for either shManb1 or shHexA are less than 0.5, indicating significant beneficial effects on fiber size by these two shRNAs, while the effects by shHexA are milder than shMan2b1. Such effects can be simply missed by traditional measurements of median, average, and Student’s *t*-test ^38^.

**Table 1.**
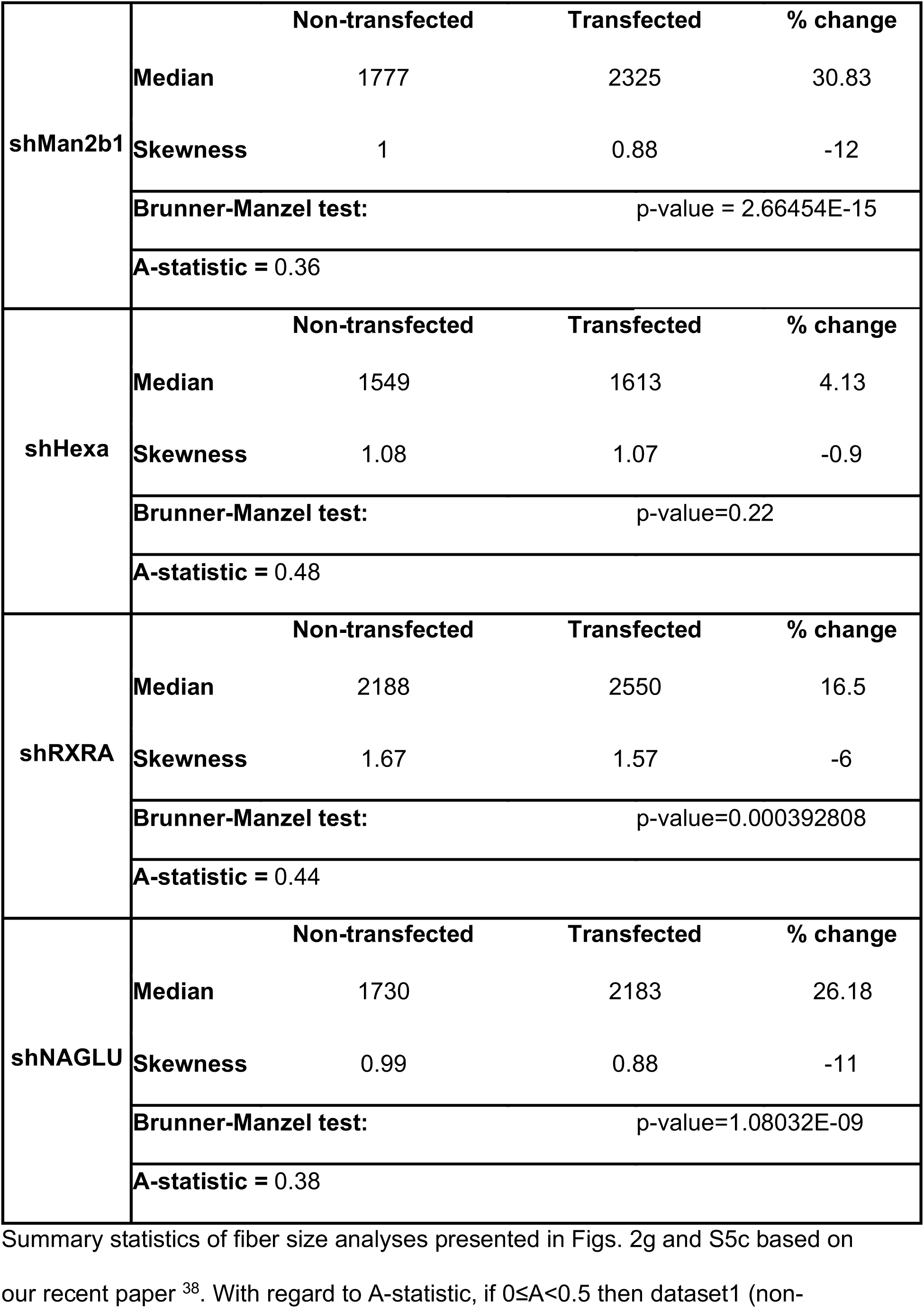
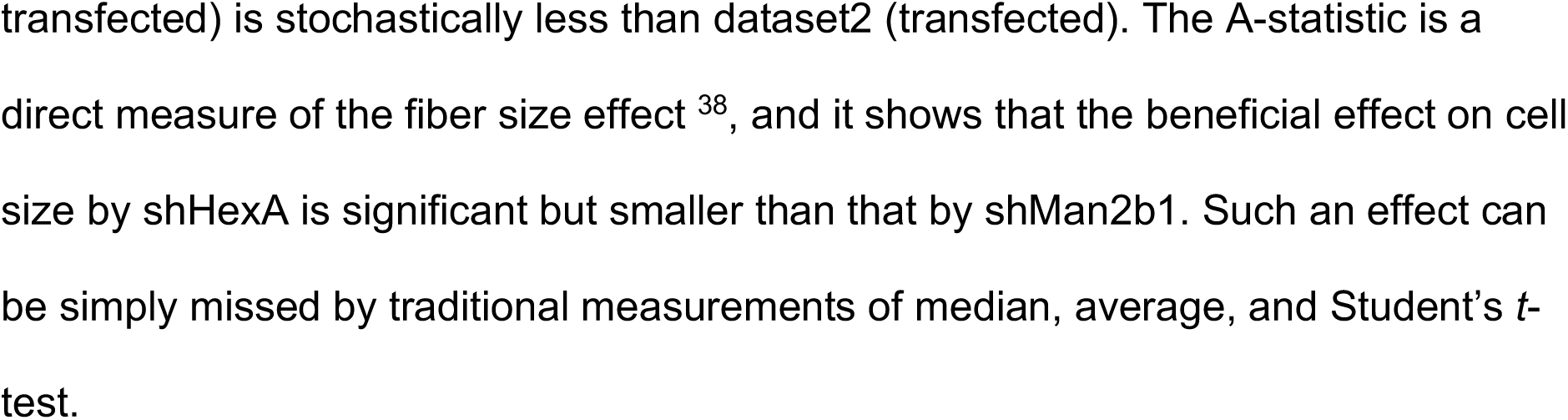

The accumulation of insulin receptor on the muscle membrane (Fig. 2d) together with the enhanced insulin-PI3K-AKT signaling (Fig. 2e) can account for the attenuation of fiber atrophy during fasting (Fig. 2g). Interestingly, the beneficial effects on fiber size, β-dystroglycan glycosylation, and β-dystroglycan-insulin receptor-plakoglobin association by shMan2b1 appeared to be greater than the ones observed by shHexA, suggesting that Man2b1 has a more prominent role in mediating β-dystroglycan deglycosylation during fasting. These beneficial effects on muscles by shMan2b1 and shHexA did not result from a shift from slow-to-fast fibers because the mean number of *MYH7* expressing slow fibers per muscle was similar to shLacz expressing muscles (6.09%±0.73 in shLacz, 6.42%±0.20 in shMan2b1, and 5.70%±0.43 in shHexA expressing muscles, n=2)(Fig. S1d).

In line with these observations, Nycodenz flotation gradients of crude lysosomal extracts indicated that in the muscles expressing shMan2b1 or shHexA, much less of β-dystroglycan, insulin receptor, and plakoglobin were recovered in lysosomes compared with atrophying muscles expressing control shRNA (shLacz)(Figs. 2h and S1e). In these muscles, β-dystroglycan-insulin receptor-plakoglobin assemblies were stabilized on the muscle membrane (Fig. 2d-e), when their trafficking to lysosomes appears to be reduced (Fig. 2h). The accumulation of glycosylated β-dystroglycan on the membrane of muscles expressing shMan2b1 or shHexA suggested that β-dystroglycan deglycosylation precedes internalization and degradation in the lysosome. Because the glycan chains of membrane proteins usually protrude into the extracellular space, we hypothesized that Man2b1 and HexA are secreted into the extracellular milieu, as has been suggested in dendritic cells ^39^, from where they could act on β-dystroglycan to deglycosylate it. In fact, immunofluorescence staining of muscle cross sections using Man2b1 antibody (the HexA antibody was not suitable for immunofluorescence) and super-resolution Structured Illumination Microscopy (SIM) revealed that at 1 d of fasting, Man2b1 was secreted into the extracellular space (extracellular areas are marked with a broken line) and showed a punctate distribution (indicative of its presence in vesicles) along the muscle membrane in close proximity to β-dystroglycan (Fig. S2a). Its levels in the extracellular space increase already 1 d after food deprivation, just before internalization of insulin receptor-β-dystroglycan is rapid, but not if muscle fibers were transfected with shManb1 (Fig. S2a). Secretion of Man2b1 does not seem to involve secretory autophagy because the Man2b1 positive extracellular foci were not enriched with LC3 (Fig. S2b). At 2 d of fasting, when DGC destabilization and internalization by the autophagy machinery is accelerated (Fig. 1c-d), Man2b1 was mostly intracellular (Fig. S2a). It is noteworthy that the overall increase in Man2b1 staining intensity during fasting compared with fed control is in line with its induction (mRNA and protein, see Figs. 2b-c and 4e). In addition, Man2b1, Hexa and β-dystroglycan were associated with each other during fasting because they co-precipitated with β-dystroglycan from membrane extracts (Fig. S2c).

**Figure 4.**
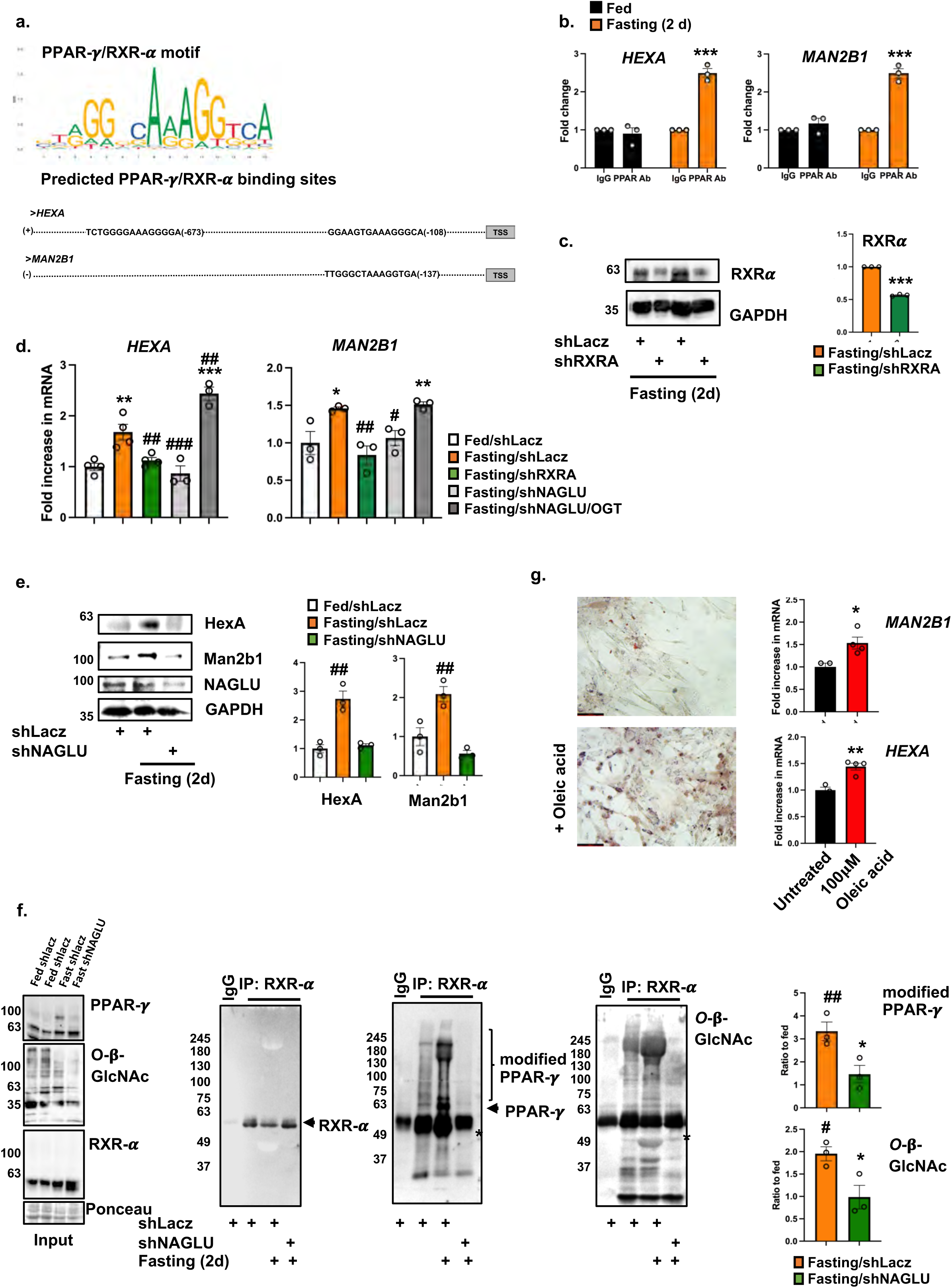
NAGLU mediates PPAR-γ/RXR-α complex formation and the resulting induction of *MAN2B1* and *HEXA*. (a) Top: PPAR-γ/RXR-α binding motif was obtained with FIMO tool. Bottom: predicted PPAR-γ/RXR-α binding sites in the promoter regions of the genes *MAN2B1* and *HEXA*, and their distances from TSS. (b) PPAR-γ binds the promoter regions of *MAN2B1* and *HEXA* genes. ChIP was performed on muscles from fed or fasted mice using PPAR-γ antibody or non-specific IgG control. Data is plotted as fold change relative to IgG control. Mean ± SEM, n = 3 mice per condition. ***, P < 0.0005 *vs.* IgG and PPAR-γ antibody in fed, by two-way ANOVA. The data are representative of three independent experiments. (c) Muscle homogenates were analyzed by immunoblotting. Ponceau S staining is shown as a loading control for RXR-α blot. The data are representative of three independent experiments. Graph depicts densitometric measurements of RXR-α blots as mean fold change relative to shLacz ± SEM. n=3 mice. ***, p<0.0005 *vs.* shLacz by one-tailed paired Student’s *t* test. (d) RT-PCR of mRNA preparations from normal or atrophying muscles expressing shLacz, shRXRA, or shNAGLU, or co-expressing shNAGLU and Myc-OGT using specific primers for *MAN2B1* and *HEXA*. Data are plotted as the mean fold change relative to control ± SEM. n=3-4 mice per condition. *, p<0.05; **, p<0.005; ***, p<0.0005 *vs.* shLacz in fed and #, p<0.05; ##, p<0.005; ###, p < 0.0005 *vs.* shLacz in fasting by ANOVA. (e) Whole cell extracts of muscles expressing shLacz or shNAGLU from fed or fasted mice were analyzed by immunoblotting. The data are representative of three independent experiments. Data are plotted as the mean fold change relative to fed ± SEM. n=3 mice per condition. ##, p<0.005 *vs.* fed and shNAGLU in fasting by ANOVA. (f) The enhanced PPAR-γ-RXR-α association during fasting is completely blocked when NAGLU is downregulated. RXR-α was immunoprecipitated from equal amounts of whole cell extracts from muscles expressing shNAGLU or shLacz from fed or fasted mice. Protein precipitates were analyzed by immunoblotting using antibodies against PPAR-γ, RXR-α, and *O*-β-GlcNAcylation. Asterisks represent nonspecific bands. The data are representative of three independent experiments. Data are plotted as the mean fold change relative to fed ± SEM. n=3 mice per condition. #, p<0.05: ##, p < 0.005 *vs.* fed; *, p < 0.05 vs. shLacz in fasting by ANOVA. (g) C2C12 cells were treated with oleic acid or left untreated, and then stained with Oil Red O or subjected to RT-PCR using specific primers to *HEXA* and *MAN2B1*. The data are representative of two independent experiments. Data are plotted as the mean fold change relative to untreated control ± SEM. n=4 wells of cells. *, p<0.05; **, p<0.005 *vs.* untreated by one-tailed unpaired Student’s *t* test.

Surprisingly, the downregulation of NAGLU with a specific shRNA (shNAGLU) also caused a marked reduction in the lysosomal content of β-dystroglycan, insulin receptor, and plakoglobin (Figs. 2h and S1e). This is surprising because NAGLU specifically removes GlcNAc units in α-linkage, and not the type of glycosylation on β-dystroglycan (N-linked)(Fig. 2a). Thus, the effect of NAGLU downregulation on β-dystroglycan-insulin receptor-plakoglobin turnover is probably indirect (see below). It is noteworthy that while NAGLU deficiency through development, in human ^40^ and in knockout mice ^41^, causes accumulation of hepran sulphate in lysosomes and impairs autophagy, the transient selective downregulation of NAGLU in wild-type fully developed mouse muscle by the transfection of shNAGLU does not impair autophagy (Fig. S3a) nor does it cause heparan sulphate accumulation (Fig. S3b), indicating proper lysosome function. Therefore, in muscles of fasted mice, the transient downregulation of NAGLU selectively inhibited β-dystroglycan-insulin receptor-plakoglobin trafficking to lysosomes. The shNAGLU seemed to rather slightly enhance autophagy flux (shown by LC3-II/LC3-I ratio in Fig. S3a), but this effect must be minor because it did not cause accumulation of autophagosomes (Fig. 2h, compare shNAGLU with shLacz).

### β-Dystroglycan is deglycosylated by a mechanism requiring NAGLU

To determine whether NAGLU in fact mediates β-dystroglycan deglycosylation, we initially performed immunofluorescence staining of muscle cross sections with β-dystroglycan and NAGLU antibodies. NAGLU is known to be a lysosomal enzyme, and accordingly it colocalizes with LAMP1 ^40^, also in skeletal muscle (Fig. 3a). As shown in Fig. 3b and our prior investigations ^6^, the membrane content of β-dystroglycan was reduced 2 d after food deprivation, likely because this protein was shunted to destruction in lysosomes (Fig. 1c-d). Accordingly, the colocalization of β-dystroglycan with the lysosomal enzyme NAGLU increased during fasting (Fig 3b), when the mRNA and protein levels of NAGLU were also elevated (Figs. 2b and 3a-b). To learn whether NAGLU influences β-dystroglycan glycosylation and thus β-dystroglycan-insulin receptor content on the muscle membrane, we electroporated shNAGLU into TA muscles, which efficiently downregulated NAGLU (Figs. 3c and S3b). During fasting, the membrane content of glycosylated β-dystroglycan, as well as insulin receptor and plakoglobin was reduced, but not if NAGLU was downregulated with shNAGLU (Fig. 3d). Whether or not NAGLU was downregulated, the membrane content of the DGC component α--1-Syntrophin did not change markedly during fasting from that in the fed control (Fig. 3d). The deglycosylation of β-dystroglycan during fasting was specifically mediated by NAGLU because the increase in the amount of glycosylated β-dystroglycan in muscles expressing shNAGLU was completely reversed by the ectopic co-expression of Myc-tagged full length NAGLU (Fig. S3c).

**Figure 3.**
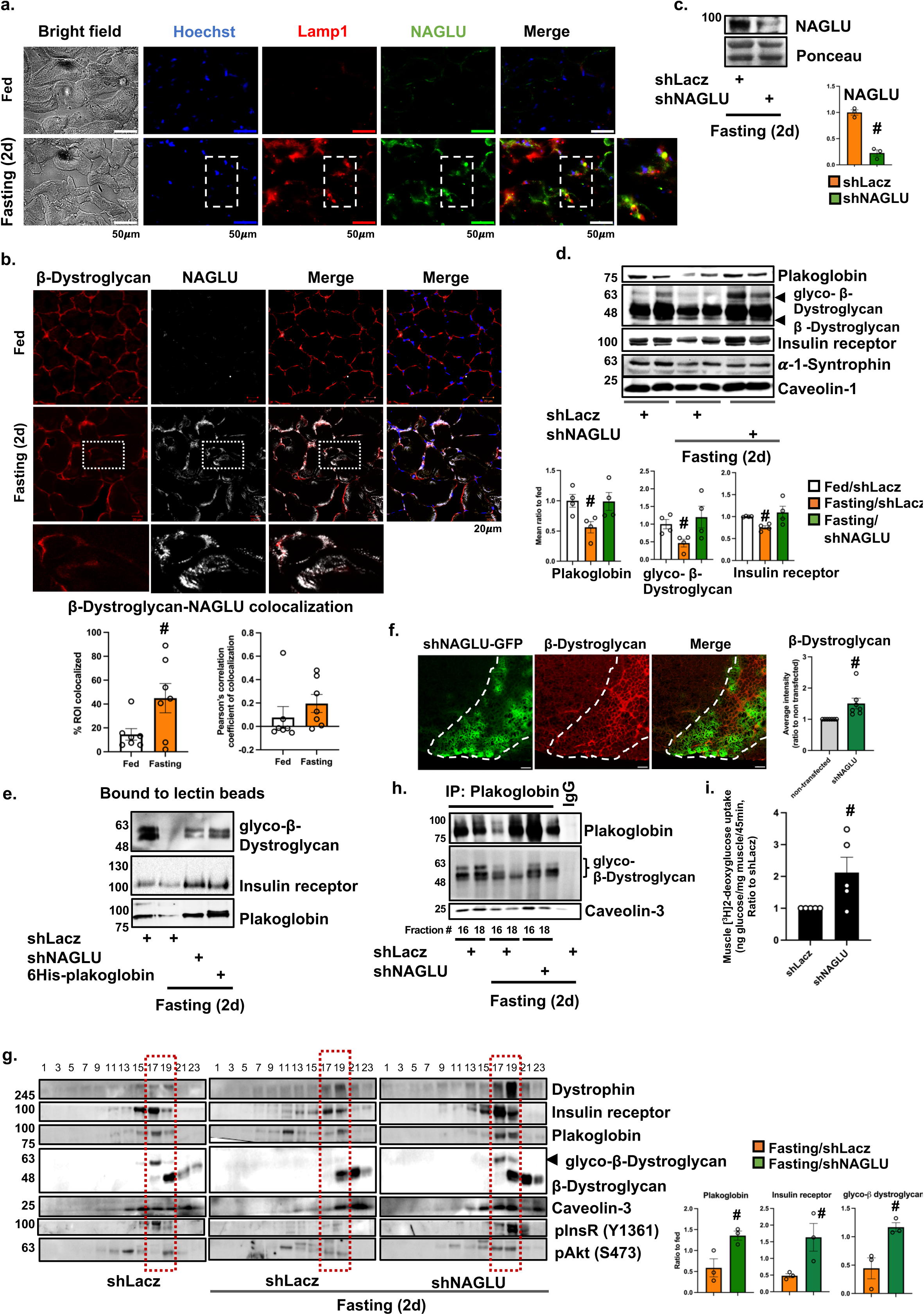
β-Dystroglycan is deglycosylated by a mechanism requiring NAGLU. (a) Muscle cross-sections from fed or fasted mice were stained with lamp1 and NAGLU antibodies. Inserts show areas of lamp1-NAGLU colocalization. Far left images are transmitted light. Bar, 50 μm. The data are representative of two independent experiments. (b) Muscle cross-sections from fed or fasted mice were stained with anti-β-dystroglycan and anti-NAGLU. Inserts show areas of β-dystroglycan-NAGLU colocalization. Bar, 20 μm. Graphs depict colocalization of indicated proteins in region of interest (ROI, region of co-occurrence), and the corresponding Pearson’s correlation coefficients of colocalization. The data are representative of 3 mice per condition and three independent experiments. Means are depicted ± SEM. *, p<0.05 *vs.* fed by one-tailed unpaired Student’s *t* test. (c) Whole cell extracts were analyzed by immunoblotting. Ponceau S staining is shown as a loading control for NAGLU blot. The data are representative of three independent experiments. Graph depicts densitometric measurement of NAGLU blots as ratio to shLacz (mean ± SEM). n=3 mice per condition. #, p<0.05 *vs.* shLacz by one-tailed paired Student’s *t* test. (d-e) During fasting, expression of shNAGLU causes accumulation of glycosylated β-dystroglycan, insulin receptor, and plakoglobin on the muscle membrane. (d) Equal fractions of membrane extracts from muscles expressing shNAGLU or shLacz from fed or fasted mice were analyzed by immunoblotting. Graphs depict mean ratio to fed ± SEM. n=4 mice per condition. #, p<0.05 *vs.* fasting shNAGLU by ANOVA. (e) Membrane extracts from whole TA muscles expressing shLacz, shNAGLU or 6His-plakoglobin from fed or fasted mice were subjected to affinity purification of glycoproteins using SNA Lectin-coated beads. Protein eluates were analyzed by immunoblotting. The data are representative of two independent experiments. (f) During fasting, expression of shNAGLU markedly attenuates β-dystroglycan loss. Cross-section of TA muscle expressing shNAGLU (green) from fasted mice was stained with β-dystroglycan antibody (red). β-dystroglycan is lost in fibers that do not express shNAGLU. Bar, 50 μm. The data are representative of three independent experiments. Graph depicts average intensity of β-dystroglycan in non-transfected *vs*. shNAGLU transfected fibers in the same muscle. mean ± SEM. n=3 mice. #, p<0.05 *vs*. non-transfected fibers. (g-h) Membrane extracts were isolated from whole TA muscles expressing shNAGLU or shLacz from fed or fasted mice, and were analyzed by glycerol gradient fractionations (g) followed by plakoglobin immunoprecipitation (h) and immunoblotting. The data are representative of three independent experiments. Graphs in (g) depict densitometric measurements as mean ratio to fed ± SEM. n=3 mice per condition. #, p<0.05 *vs.* fasting shLacz by one-tailed unpaired Student’s *t* test. (i) In vivo [^3^H]-2-deoxyglucose uptake by muscles expressing shLacz or shNAGLU (contralateral limbs) during fasting (2 d) is plotted as mean ratio to shLacz control ± SEM. Data represents ng glucose/mg muscle/45 min. Mean ± SEM, n = 5 mice. #, P < 0.05 *vs.* shLacz by one-tailed paired Student’s *t* test. The data are representative of two independent experiments.

To confirm these findings by an independent approach, we analyzed more directly the effect of NAGLU downregulation on the total membrane content of glycosylated-β-dystroglycan upon fasting. Using SNA Lectin-coated beads we isolated glycosylated proteins from appropriately solubilized membrane extracts from whole muscles expressing shNAGLU or shLacz, and analyzed protein eluates by SDS-PAGE and immunoblotting (Fig. 3e). The SNA lectin-based affinity purification relies on the high and preferential affinity of the SNA lectin to sialic acid, which is terminally attached to N-linked and O-linked glycoproteins. Glycosylated β-dystroglycan was efficiently purified with SNA lectin beads from normal muscle (Fig. 3e), where β-dystroglycan is present on the plasma membrane in its glycosylated form (Figs. 1a and 3d). The membrane extracts isolated from atrophying muscles contained considerably lower amounts of glycosylated-β-dystroglycan (Figs. 1a and 3d), and therefore β-dystroglycan could not be retrieved with the lectin-based affinity purification approach (Fig. 3e). By contrast, glycosylated β-dystroglycan was efficiently purified with SNA lectin beads from membrane extracts of the atrophying muscles expressing shNAGLU (Fig. 3e), because in these muscles β-dystroglycan accumulated in its glycosylated form (Fig. 3d). Interestingly, insulin receptor and plakoglobin were co-purified with SNA-coated lectin beads mostly from membrane extracts containing glycosylated-β-dystroglycan (i.e. normal muscle, and atrophying muscle expressing shNAGLU); their affinity purification was probably indirect and resulted from association with glycosylated-β-dystroglycan (Fig. 3e). These findings were further validated by the analysis of muscles expressing 6His-plakoglobin. Previously, we demonstrated that plakoglobin accumulation prevents β-dystroglycan deglycosylation and protects β-dystroglycan-insulin receptor co-assembly from LC3-mediated degradation ^6^. Accordingly, incubation of membrane extracts from muscles expressing 6His-plakoglobin with lectin beads resulted in efficient purification of glycosylated β-dystroglycan and the attached insulin receptor and plakoglobin (Fig. 3e). Thus, during fasting, NAGLU mediates β-dystroglycan deglycosylation.

To determine whether this deglycosylation leads to degradation, we performed immunofluorescence staining of atrophying TA muscle expressing shNAGLU from fasted mice using an antibody against β-dystroglycan. In non-transfected fibers, where β-dystroglycan is lost, the staining intensity of β-dystroglycan was low (Fig. 3f). However, transfection of muscle fibers with shNAGLU (also express GFP) markedly attenuated this loss of β-dystroglycan, which accumulated on the muscle membrane (Fig. 3f) in its glycosylated form (Fig. 3d-e). In fact, glycerol gradient fractionation of membrane extracts from muscles expressing shLacz or shNAGLU from fed or fasted mice indicated that the membrane content of glycosylated-β-dystroglycan-insulin receptor-plakoglobin assemblies is reduced during fasting, but not when NAGLU was downregulated (Fig. 3g). This attenuation in β-dystroglycan loss in atrophying muscles lacking NAGLU (Fig. 3f-g) led to stabilization of glycosylated-β-dystroglycan-plakoglobin co-assemblies on the muscle membrane (Fig. 3g), where plakoglobin is expected to protect β-dystroglycan from LC3-mediated loss (Fig. 3e and ^6^). Accordingly, immunoprecipitation of plakoglobin from the glycerol gradient fractions (fractions #16 and 18 in Fig. 3g) demonstrated that plakoglobin-glycosylated-β-dystroglycan association was reduced in fasting, but not when NAGLU was downregulated (Fig. 3h). Moreover, the stabilization of this co-assembly on the muscle membrane was sufficient to enhance insulin signaling (Fig. 3g) and glucose uptake (Fig. 3i). Thus, NAGLU appears critical for β-dystroglycan deglycosylation and β-dystroglycan-insulin receptor loss during fasting.

### NAGLU promotes PPAR-γ/RXR-α complex activity, which induces *MAN2B1* and ***HEXA*.**

We hypothesized that NAGLU indirectly promotes β-dystroglycan deglycosylation by activating the transcription factor responsible for *MAN2B1* and *HEXA* induction. To test this idea and identify the transcription factor responsible for their induction, we initially searched for potential transcription factor-binding sites in the promoter regions of these genes using the CISTROME ChIP-Seq portal ^42^. Several potential transcription factor-binding motifs were identified for the two enzymes (Table S1), while only binding motifs for the transcription factor RXR-α were predicted by CISTROME in both *MAN2B1* and *HEXA* genes. Induction of RXR-α target genes requires heterodimerization of RXR-α with the nuclear hormone receptor, peroxisome proliferator-activated receptor (PPAR), to form a functional transcription factor complex ^43^. Among the three known PPAR isoforms, PPAR-α, PPAR-β/δ and PPAR-γ, PPAR-γ is considered a master regulator of the physiological systemic response to insulin, especially in fat, liver and muscle ^44^, and activates physiological survival mechanisms in low glucose states in humans ^45^. Therefore, we investigated whether this nuclear receptor contributes to the induction of *MAN2B1* and *HEXA* under low glucose and low insulin conditions (i.e. during fasting) by forming a functional transcriptional heterodimer with RXR-α. Initially, it was important to confirm that PPAR-γ/RXR-α binding motif is in fact present in the promoter regions for both genes. For this purpose, we extracted PPAR-γ/RXR-α binding motif using FIMO tool (http://meme-suite.org/doc/fimo.html) (Fig. 4a) and used it as input to scan the promoter regions of human *MAN2B1* and *HEXA* genes at 10kb from Transcription Start Site (TSS). As shown in Fig. 4a, binding motifs for the transcription factor PPAR-γ/RXR-α were predicted by FIMO in both *MAN2B1* and *HEXA* genes. To confirm that PPAR-γ/RXR-α binds the promoter regions of these genes, we performed chromatin immunoprecipitation (ChIP) from muscles of fed or fasted (2 d) mice using a PPAR-γ antibody or a non-specific IgG as control. RT-PCR analysis of immunoprecipitates using specific primers for binding motifs within mouse *MAN2B1* and *HEXA* promoters (Fig. 4a and Table S2) confirmed enhanced binding of PPAR-γ to these genes during fasting but not in fed conditions or in control samples containing a non-specific IgG (Fig. 4b).

To test whether RXR-α is in fact essential for the induction of these genes during fasting, we suppressed RXR-α expression by electroporation into mouse TA of shRNA plasmid (shRXRA), which efficiently reduced RXR-α protein levels in atrophying muscles below the levels in shLacz expressing controls (Fig. 4c). As shown above (Fig. 2c), *MAN2B1* and *HEXA* were induced in atrophying muscles during fasting (Fig. 4d). However, the downregulation of RXR-α resulted in a significant decrease in their expression, and their mRNA levels no longer differed from those in fed controls (Fig. 4d). Thus, during fasting, RXR-α is responsible for Man2b1 and HexA induction.

Intriguingly, the downregulation of NAGLU with shNAGLU also resulted in a marked decrease in the levels of Man2b1 and HexA mRNA (Fig. 4d) and protein (Fig. 4e), indicating that NAGLU is required for RXR-α-mediated *MAN2B1* and *HEXA* induction. To determine if NAGLU facilitates β-dystroglycan deglycosylation and loss by promoting RXR-α-mediated *MAN2B1* and *HEXA* expression, we investigated whether the activity of RXR-α during fasting, i.e. the formation of a functional transcriptional heterodimer with PPAR-γ, is affected by the levels of NAGLU. PPAR-γ could be coprecipitated with RXR-α from normal muscles, and fasting elicited a remarkably greater association (Fig. 4f). This change in PPAR-γ association seemed to require NAGLU because downregulation of NAGLU during fasting prevented the association of PPAR-γ with RXR-α (Fig. 4f). Thus, during fasting, NAGLU facilitates PPAR-γ and RXR-α association. Before association with RXR-α, PPAR-γ may be activated in fasting by free fatty acids or their metabolites released from lipid reserves^46^. Activation of PPAR-γ likely involves association with oleic acid, whose circulating blood levels increase during fasting ^47^, and which can induce PPAR-γ target genes, *MAN2B1* and *HEXA*, in cultured C2C12 myotubes (Fig. 4g, red lipid staining confirms penetration of oleic acid into the cells).

Previously, the activity of PPAR-γ has been shown to also be regulated by *O*-GlcNAcylation ^48^. The *O*-GlcNAc posttranslational modification is often involved in regulation of cytosolic and nuclear proteins function, including transcription factors ^49^. To determine if PPAR-γ is *O*-GlcNAcylated during fasting, we analyzed the RXR-α coprecipitates (Fig. 4f) with *O*-GlcNAcylation specific antibody. In normal muscle, *O*-GlcNAcylated species of PPAR-γ bound RXR-α (Fig. 4f). This modification of PPAR-γ probably occurs at multiple sites to regulate its activity, as had been proposed also for other transcription factors ^48,49^. In fact, upon fasting, when PPAR-γ association with RXR-α increased (Fig. 4f), PPAR-γ within PPAR-γ/RXR-α complex was heavily modified (Fig. 4f) and stimulated the induction of *MAN2B1* and *HEXA* (Fig. 4d). Interestingly, RXR-α within this complex was not modified by *O*-GlcNAcylation, when its association with *O*-GlcNAcylated PPAR-γ required NAGLU (Fig. 4f). Because protein *O*-GlcNAcylation results in small changes in protein molecular weight, the high molecular weight species of *O*-GlcNAcylated PPAR-γ probably represent an additional modification ^50^. Accordingly, PPAR-γ is also SUMOylated by SUMO1 during fasting (Fig. S4), a modification that is known to promote PPAR-γ stabilization ^51^.

### NAGLU mediates PPAR-γ O-GlcNAcylation by promoting OGT stabilization

These findings suggested that during fasting NAGLU promotes PPAR-γ *O*-GlcNAcylation and hence PPAR-γ/RXR-α complex activity. Because the primary enzyme responsible for protein *O*-GlcNAcylation is *O*-GlcNAc transferase (OGT) ^49^, we determined whether NAGLU influences its levels in muscle during fasting. Analysis of muscle homogenates by immunoblotting using OGT antibody showed that OGT protein levels dramatically decreased when NAGLU was downregulated, indicating that NAGLU is essential for OGT stability during fasting (Fig. 5a). This stabilization results from reduced degradation, and not from increased gene expression, because during fasting *OGT* is not induced (Fig. 5b). Consistently, in the atrophying muscles expressing shNAGLU, the total levels of *O*-GlcNAcylated proteins were markedly reduced compared to muscles expressing shLacz (Fig. 5a). Moreover, in the atrophying muscles expressing shNAGLU, transfection of Myc-tagged full-length OGT completely reversed the inhibitory effects imposed by NAGLU downregulation and promoted *MAN2B1* and *HEXA* induction (Fig. 4d). Thus, the NAGLU-dependent stabilization of OGT appears critical in causing PPAR-γ *O*-GlcNAcylation, and consequently PPAR-γ/RXR-α-mediated *MAN2B1* and *HEXA* induction. Immunoprecipitation of heparan sulfate from muscle homogenates using a specific antibody indicated that OGT bound heparan sulfate only in atrophying muscles expressing shNAGLU but not in control muscles (expressing shLacz)(Fig. 5c). These findings suggest that association of OGT with heparan sulfate promotes OGT loss, and during fasting NAGLU enhances OGT stabilization by promoting heparan sulfate degradation.

**Figure 5.**
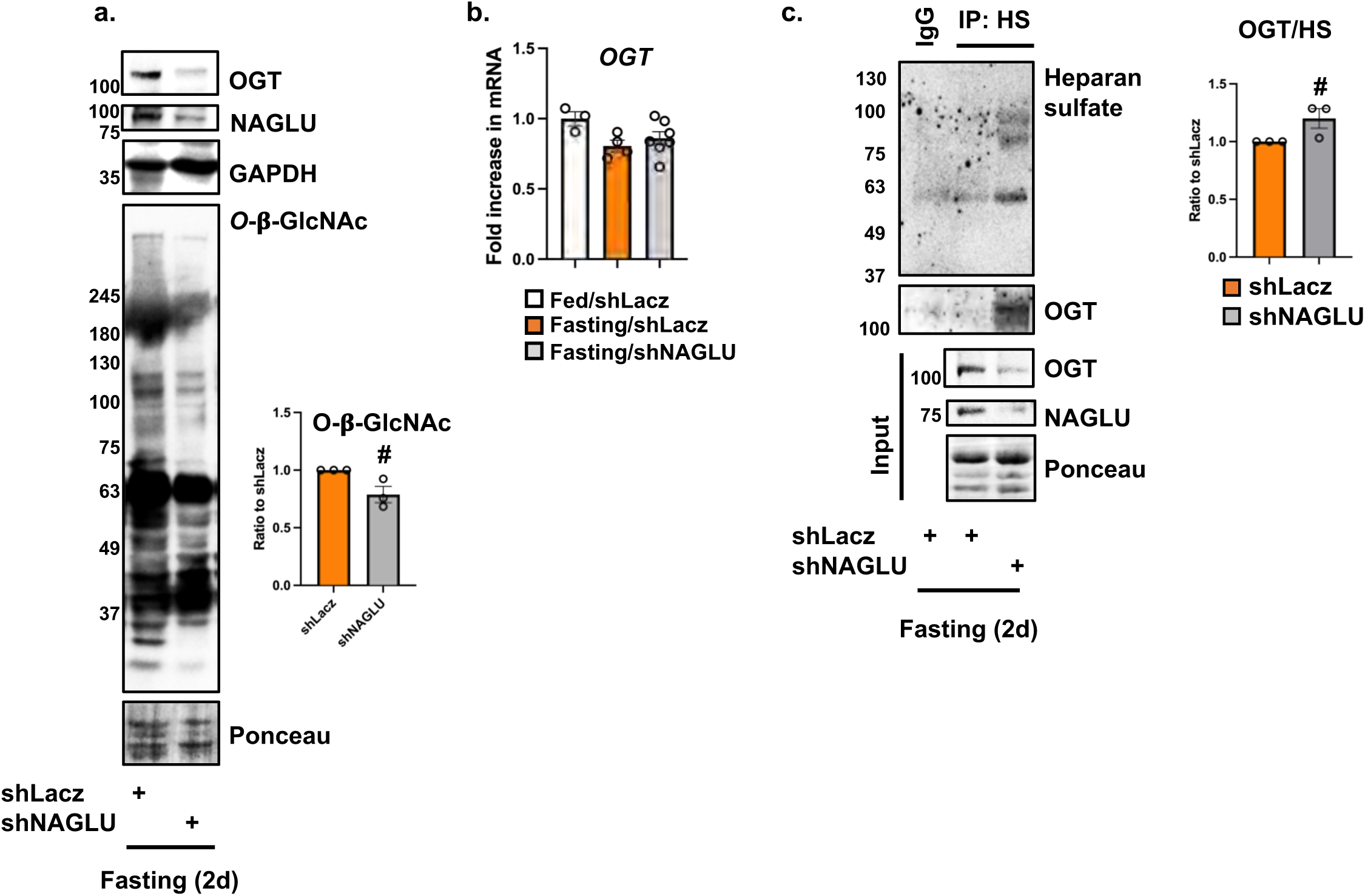
NAGLU mediates PPAR-γ O-GlcNAcylation by promoting OGT stabilization. (a) NAGLU in essential for OGT stability during fasting. Homogenates of muscles expressing shLacz or shNAGLU from fasted mice were analyzed by immunoblotting. Protein *O*-β-GlcNAcylation was detected using an *O*-β-GlcNAc antibody. Ponceau S staining is shown as a loading control for *O*-β-GlcNA blot. The data are representative of three independent experiments. Graph depicts densitometric measurements of *O*-β-GlcNA blots as mean fold change relative to shLacz ± SEM. n=3 mice per condition. #, p<0.05 *vs.* shLacz by one-tailed paired Student’s *t* test. (b) *OGT* is not induced in muscle during fasting. RT-PCR of mRNA preparations from normal or atrophying muscles expressing shLacz or shNAGLU using specific primers for *OGT*. Data are plotted as the mean fold change relative to fed control ± SEM. n=3-7 mice per condition. Not significant by ANOVA. (c) Heparan sulfate (HS) was immunoprecipitated from muscles expressing shLacz or shNAGLU from fasted mice, and protein precipitates were analyzed by immunoblotting using the indicated antibodies. Ponceau S staining is shown as a loading control for input blots. The data are representative of three independent experiments. Graph depicts densitometric measurements as mean OGT/HS ratio relative to shLacz ± SEM. n=3 mice per condition. #, p<0.05 *vs.* shLacz by one-tailed unpaired Student’s *t* test.

Further studies determined the sequential effects on β-dystroglycan deglycosylation and β-dystroglycan-insulin receptor loss. In muscles expressing shRXRA, β-dystroglycan, insulin receptor, and plakoglobin accumulated on the muscle membrane as intact co-assemblies (Fig. S5a). This increase in the amount of glycosylated β-dystroglycan in muscles expressing shRXRA was completely reversed by the ectopic co-expression of full-length RXR-α (Fig. S5b), indicating that the deglycosylation of β-dystroglycan during fasting is specifically mediated by RXR-α.

Moreover, these beneficial effects on β-dystroglycan glycosylation and β-dystroglycan-insulin receptor-plakoglobin association also attenuated fiber atrophy when RXR-α or NAGLU were downregulated (Fig. S5c and Table 1). These beneficial effects on fiber size by shRXRA or shNAGLU did not result from a shift from slow-to-fast fibers because the mean number of *MYH7* expressing slow fibers per muscle transfected with shRXRA or shNAGLU was similar to shLacz expressing controls (6.09%±0.73 in shLacz, 5.79%±0.27 in shRXRA, and 5.86%±0.33 in shNAGLU expressing muscles, n=2)(Fig. S1d). Thus, during fasting, NAGLU function is critical in facilitating PPAR-γ/RXR-α activity, which triggers the induction of *MAN2B1* and *HEXA*, the glycoside hydrolyses that mediate β-dystroglycan deglycosylation and the resulting loss of the insulin receptor.

## DISCUSSION

These studies identified β-dystroglycan deglycosylation as a key molecular event in the destabilization of insulin receptor during prolonged starvation. We recently showed that DGC-insulin receptor clusters are present on the muscle membrane and are dynamically regulated by autophagy ^6^, and the present studies demonstrate that β-dystroglycan deglycosylation is a critical step in promoting insulin receptor loss. Therefore, the physiological response of skeletal muscle to fasting, which involves reduced transmission of insulin-mediated growth signals from the cell surface, occurs not only through the reduced binding of insulin to its receptor, but also via enhanced internalization and loss of the insulin receptor, which appears to be dependent on β-dystroglycan deglycosylation.

The conserved post-translational modification of N-linked glycosylation occurs by the attachment of glycan chains to asparagine residues of polypeptides in the endoplasmic reticulum (ER). While the common knowledge is that N-linked glycosylation is primarily required for trafficking to the plasma membrane, we discover here that this type of glycosylation has an additional functional role in maintenance of tissue homeostasis and response to insulin. Glycosylation of β-dystroglycan is known to be required not only for trafficking to the plasma membrane but also for maintenance of DGC integrity ^16^, and as shown here, fasting elicits a reduction in glycosylated β-dystroglycan on the muscle membrane and a simultaneous internalization and destruction of β-dystroglycan-insulin receptor-plakoglobin co-assemblies. It is possible, that trafficking of these proteins from the ER to the plasma membrane is also reduced during fasting by a mechanism requiring Man2b1 and HexA, although our crude lysosomal extracts contained only small traces of ER and Golgi (Fig. 1c). In normal muscle, β-dystroglycan is N-linked glycosylated (Fig. 2a), as reported in other tissues ^16,22,36^, and its deglycosylation during fasting is mediated by the lysosomal glycoside hydrolyses Man2b1 and HexA (Fig. 2). Because these enzymes are exoglycosidases, they should act on β-dystroglycan to deglycosylate it only after certain residues, such as sialic acid (Fig. 3e), have been removed by other enzymes; hence, their effects on β-dystroglycan are probably indirect.

Our present studies and prior ones ^6^ clearly demonstrate that during fasting, both unmodified and glycosylated forms of β-dystroglycan are shunted to lysosomes (with preference to the unmodified deglycosylated form), where they are degraded together with the insulin receptor. Thus, β-dystroglycan is not only deglycosylated but is also degraded. Accordingly, during fasting, unmodified and glycosylated β-dystroglycan accumulate in autophagosomes on colchicine treatment (Fig. 1d). In addition, inhibition of β-dystroglycan deglycosylation (as shown here by Man2b1, HexA or NAGLU downregulation) reduces trafficking to lysosomes (Fig. 2e-h), leading to stabilization of both forms of β-dystroglycan (i.e. unmodified and glycosylated) on the plasma membrane. How Man2b1 and HexA promote trafficking to lysosomes is still unclear. Possibly, these enzymes are secreted into the extracellular milieu via exosomes, as has been suggested in dendritic cells ^39^ and as we show here for Man2b1 (Fig. S2), where they may promote β-dystroglycan deglycosylation, consequently facilitating DGC destabilization and internalization by the autophagy machinery. Its secretion does not seem to occur via secretory autophagy because Man2b1 extracellular puncta were not enriched with the bona fide autophagy marker, LC3. Clearly, deglycosylation of all β-dystroglycan molecules on the muscle membrane is not required to promote trafficking to lysosomes during fasting, because some fraction of glycosylated β-dystroglycan can also be detected in lysosomes (Fig. 1c). Glycosylated β-dystroglycan is likely dragged into internalization alongside deglycosylated β-dystroglycan within the formed vesicles.

To our knowledge, the functions of Man2b1 and HexA in muscle atrophy have not been investigated previously, and we show they are of prime importance in promoting β-dystroglycan deglycosylation and the coupled loss of insulin receptor. These exoglycosidases catalyze the breakdown of complex N-linked glycans ^52^, and may serve similar regulatory roles in controlling DGC integrity and growth of other tissues because Man2b1 and HexA are expressed in all cells. Recent studies in kidney cells demonstrated that mutations preventing N-linked glycosylation of membrane proteins cause a reduction in the membrane content of insulin receptors ^53^. These effects can now be explained in part based on our findings that a reduction in the membrane content of N-linked glycosylated β-dystroglycan is coupled to insulin receptor loss. Interestingly, early studies demonstrated that insulin receptors are N-linked and O-linked glycosylated in fat and brain ^54,55^, although the effects of these posttranslational modifications on insulin receptor function or stability remain uncertain. When associated with N-linked glycosylated β-dystroglycan on skeletal muscle membrane, insulin receptor appears mainly in its native form (Fig. 3e), and the formed co-assembly is an important signaling hub regulating cell growth ^6^.

These studies also identified a critical role for the lysosomal enzyme NAGLU, which is induced in fasting and promotes β-dystroglycan deglycosylation and the coupled insulin receptor loss. These effects are not caused by NAGLU directly deglycosylating β-dystroglycan, as NAGLU specifically processes GlcNAc units that are terminally attached to glycan chains in α-linkage, while β-dystroglycan is N-linked glycosylated in β-linkage. Instead, NAGLU mediates β-dystroglycan deglycosylation via Man2b1 and HexA through activation of the transcription factor responsible for their induction, i.e. PPAR-γ/RXR-α (Fig. 4). Further investigations into NAGLU-mediated β-dystroglycan deglycosylation uncovered a critical role for NAGLU in promoting OGT stability and consequently PPAR-γ *O*-GlcNAcylation and PPAR-γ/RXR-α activation during fasting. NAGLU mediates OGT stability by promoting heparan sulfate degradation, consequently liberating OGT from association with heparan sulfate. Consistently, cleavage products of heparan sulfate are known to regulate various cellular functions ^56^ through effects on protein activity and/or stability ^57^. Although NAGLU has been studied for more than four decades, the only known cellular role for this enzyme is degradation of heparan sulphate. Over 100 loss of function mutations in NAGLU gene have been identified in humans, all of which cause heparan sulphate accumulation and the lysosomal storage disease Sanfilippo syndrome type B ^40^. However, the transient selective downregulation of NAGLU in atrophying muscles from wild type mice during fasting does not inhibit heparan sulphate turnover (Fig. S3b) or lysosome function (Fig. S3a). Therefore, the accumulation of β-dystroglycan-insulin receptor-plakoglobin assemblies on the membrane in muscles expressing shNAGLU resulted from a specific effect and not from a general inhibitory effect on autophagy.

The beneficial effects of PPAR-γ activation in low energy states has been suggested before. For example, in glucose intolerant human subjects, PPAR-γ agonists can induce expression of genes that activate pathways related to fatty acids storage in lipid droplets in muscle, as a survival mechanism ^45^. Within PPAR-γ/RXR-α complex, PPAR-γ is SUMOylation by SUMO1 during fasting, when this transcription factor induces *HEXA* and *MAN2B1*. Consistently, SUMOylation by SUMO1 has been shown to be important for PPAR-γ stability and transcriptional activity ^51^. Within this complex, PPAR-γ is also modified by *O*-GlcNAcylation (Fig. 4f). This distinct form of protein glycosylation is known to regulate signaling pathways and transcription by mainly regulating protein function ^58^. It occurs in the nucleus and cytosol and involves the attachment of single GlcNAc moieties in β-linkage to serine or threonine residues on substrate proteins by the enzyme OGT ^49^. Because *O*-GlcNAc is not elongated to form more complex chains, the high molecular weight species of PPAR-γ likely indicate *O*-GlcNAcylation at multiple sites, of an already SUMOylated protein. Previously, *O*-GlcNAcylation of specific threonine 54 residue in PPAR-γ had been proposed to reduce its transcriptional activity ^48^, although mass spectrometry analysis in the same study identified several additional threonine and serine residues in PPAR-γ as *O*-GlcNAc-modified sites, whose roles in regulating PPAR-γ activity remain unknown. Whether *O*-GlcNAcylation of PPAR-γ is required for its association with RXR-α and activation is an important question for future research. In addition, although NAGLU cannot directly influence PPAR-γ *O*-GlcNAcylation, it promotes OGT stability, which is clearly essential for protein *O*-GlcNAcylation.

Dynamic recycling of insulin receptors via autophagy ^6^ or by the ubiquitin-proteasome system ^59^ serves to modulate the number of insulin receptors on the cell surface and regulate the transmission of growth signals into the cell. Perturbations to this dynamics interfere with cellular energy and metabolic homeostasis, resulting in impaired proteostasis and carbohydrate metabolism, and insulin resistance. Accordingly, hyperglycemia and hyperinsulinemia in obesity or diabetes facilitate insulin receptor recycling, which may contribute to the development of insulin resistance, and our prior ^6^ and present studies propose that these effects are linked to a reduction in glycosylated-β-dystroglycan on the muscle membrane. Therefore, accelerated insulin receptor turnover that contributes to the fall in insulin signaling during fasting, and the accompanied loss of β-dystroglycan, if persisted, may facilitate the development of insulin resistance seen in prolonged starvation ^60^. Our recent paper on the MKR type-2 diabetic mouse models demonstrated a marked reduction in insulin receptor membrane content and the bound DGC and plakoglobin ^6^. Moreover, overexpression of plakoglobin to stabilize insulin receptor on the membrane was sufficient to enhance glucose uptake in diabetic mice ^6^, indicating that a reduction in the membrane content of insulin receptor contributes to the insulin resistance in type-2 diabetes. A similar reduction in insulin receptor levels on the muscle membrane and in other tissues has been demonstrated also in diabetic human patients ^61–63^ and in mice ^64^. Because skeletal muscle is the most abundant tissue in the human body and the major site for insulin-stimulated glucose disposal, the mechanism proposed herein, of coupling β-dystroglycan deglycosylation to insulin receptor loss, probably regulates systemic energy homeostasis, linking nutrient availability to muscle growth or atrophy.

## METHODS

### Animals

All animal experiments were consistent with Israel Council on Animal Experiments guidelines and were approved by the Technion Inspection Committee on the Constitution of the Animal Experimentation. Specialized personnel provided animal care in the Institutional Animal facility. For these studies we used Hsd:ICR (CD-1) male mice at 25-30g body weight. For fasting, food was removed for cages 5 d after *in vivo* electroporation for 2 d. For colchicine treatment (Fig. 1d), mice were injected intraperitoneally with 0.4 mg/kg colchicine (Sigma C9754) or vehicle at 24 and 36 hr after food deprivation, and mice were scarified at 48 hr after food deprivation.

### *In vivo* transfection and fiber size analysis

For *in vivo* electroporation experiments, 20 μg of plasmid DNA was injected into adult mouse TA muscles, and a mild electric pulse was applied using two electrodes (12 V, 5 pulses, 200 ms intervals). Muscle transfection efficiency was estimated using a Nikon Ni-U upright fluorescence microscope with Plan Fluor 10× 0.3-NA objective lens and a Hamamatsu C8484-03 cooled CCD camera, at room temperature, and Metamorph (Molecular Devices) or Imaris (Bitplane) software ^38,65^. For rigorous biochemical analysis, muscles that are ∼ 70% transfected were used.

For fiber size analysis, cross sections of transfected muscles (from at least 4 different mice) were fixed in 4% paraformaldehyde (PFA), and fiber membrane was stained with dystrophin antibody overnight at 4° C and nuclei with Hoechst. Images were collected using a Nikon Ni-U upright fluorescence microscope with Plan Fluor 20× 0.5-NA objective lens and a Hamamatsu C8484-03 cooled CCD camera, at room temperature. Cross-sectional area of at least 500 transfected fibers (also express GFP) and the same number of adjacent non-transfected fibers in the same muscle section (20 µm) were measured using Metamorph and Imaris software ^38,65^.

### Antibodies and constructs

The plasmids encoding shRNA against Lacz, HexA, Man2b1, RXRA and NAGLU were cloned into pcDNA 6.2-GW/EmGFP-miR vector using Invitrogen’s BLOCK-iT RNAi Expression Vector Kit as before ^6,65,66^. These shRNAs were designed against the mouse genes. The Myc-OGT encoding plasmid was a generous gift from X. Yang (Yale School of Medicine, Connecticut, USA). The 6His-plakoglobin plasmid was previously described ^6^, the plasmids encoding FL-Man2b1, FL-HexA, FL-NAGLU, and FL-RXR--α- were purchased from Origene, and the plasmid encoding GFP-LC3 (Cat #24987) was from Addgene. The anti *O-*β-GlcNAc was a kind gift from G. Hart (Johns Hopkins University, Maryland, USA), and the MuRF1 antibody was previously described ^67^. Plakoglobin antibody was from Genetex (Cat# GTX15153). Anti-dystrophin (Cat# ab15277), anti-caveolin-1 (Cat# ab2910), anti-LC3B (Cat# ab48394), anti-NAGLU (Cat# ab137685), anti ATP synthase-β subunit (Cat# ab5432), and anti-lamp1 (Cat# ab24170) were from Abcam. Syntrophin (Cat# Sc50460), RXR-α- (Cat# sc46659), PPAR-γ (Cat# sc7273), calnexin (Cat# sc23954), golgin-97 (Cat# sc59820), OGT (Cat# ab74546), SQSTM1 (Cat# sc48402), HexA (Cat# sc-376777) and caveolin-3 (Cat# sc5310) antibodies were from Santa Cruz Biotechnology. The β-dystroglycan antibody was developed by Glenn E. Morris (RJAH Orthopedic hospital, Oswestry, UK, 1:100) and obtained from the Developmental Studies Hybridoma Bank, created by the NICHD of the NIH and maintained at the University of Iowa, Department of Biology, Iowa city, Iowa (Cat# MANDAG2, clone 7D11). Anti-insulin receptor (Cat# 3025), phospho-insulin receptor (Y1361)(Cat# 3023), phospho Akt S473 (Cat# 9271), and anti-Rab7 (Cat# 9367) were from cell signaling, anti-cathepsin D (Cat# MAB1029) from R&D systems, and anti-EEA (Cat# 1610456) from BD transduction laboratories. Anti-GAPDH (Cat# G8795) and MYH7 (Cat# M8421) were from Sigma, anti-Heparan sulfate (Cat# OBT1698) was from Bio-Rad, and anti Man2b1 (Cat# orb537674) from Biorbyt.

### Fractionation of mouse skeletal muscle

To obtain whole-cell extracts, TA muscles were homogenized in 19 volumes (v/w) of lysis buffer (20 mM Tris, pH 7.2, 5 mM EGTA, 100 mM KCl, 1% Triton X-100, 1 mM PMSF, 3 mM benzamidine, 10 μg/ml leupeptin, 50 mM NaF, and 2.7 mM sodium orthovanadate), and following centrifugation at 6000 × g for 20 min at 4 °C, the supernatant (i.e. whole-cell extract) was collected and stored at −80 °C.

To obtain membrane extracts, whole TA muscles were homogenized in 19 volumes (v/w) of buffer C (20 mM Tris, pH 7.6, 100 mM KCl, 5 mM EDTA, 1 mM DTT, 1 mM PMSF, 3 mM benzamidine, 10 μg/ml leupeptin, 50 mM NaF, and 1 mM sodium orthovanadate). Following centrifugation at 2,900 × g for 20 min at 4 °C, the supernatant was collected and subjected to centrifugation at 18,000 × g for 90 min at 4 °C to isolate membranes. The obtained supernatant was stored as the cytosolic fraction, and the pellet was resuspended in 100μl Buffer M (20 mM Tris, pH 7.6, 100 mM KCl, 5 mM EDTA, 1 mM DTT, 0.25% sodium deoxycholate, 1% NP-40, 1 mM sodium orthovanadate, 1 mM PMSF, 3 mM benzamidine, 10 μg/ml leupeptin, 50 mM NaF) per 50 mg TA muscle, and was rotated for 20 min at 4 °C. Following centrifugation at 100,000 × g for 30 min at 4 °C, the supernatant containing membrane extracts was stored at −80°C.

### Isolation of lysosomes by Nycodenz fractionation

To prepare a discontinuous Nycodenz gradient, 45% Nycodenz (Axis-Shield) stock solution was initially prepared in Homogenization Buffer (HB)(0.25 M sucrose, 1 mM Na2EDTA, 10 mM HEPES, pH adjusted to 7.0 with NaOH), and was used to prepare 19%, 24%, 26% and 30% Nycodenz solutions. Then, 1ml of 19% Nycodenz solution was layered at the bottom of a 5ml ultracentrifuge tube, and using a syringe, 1ml from each of the other three Nycodenz solutions (24%, 26% and 30%) were carefully underlayered in increasing density. To isolate crude lysosomal extracts, whole mouse TA muscles were homogenized in 10 volumes (v/w) of freshly made ice-cold HB. Following centrifugation at 750 × g for 10 min at 4 °C, the supernatant was retained and centrifuged at 20,000 × g for 10 min at 4 °C. The pellet was then resuspended in 150 μl HB per 50 mg TA muscle, and the obtained homogenate (crude lysosomal fraction) was mixed with 2 volumes (v/v) of 45% Nycodenz solution, and was carefully layered under the discontinuous Nycodenz gradient using a syringe. After centrifugation for 90 min at 95,000 x g and 4 °C, fractions (300 μl each) were collected from the bottom of the tube, and following TCA precipitation (10% TCA was added to samples) were analyzed by SDS-PAGE and immunoblotting.

### Glycerol gradient fractionation

Membrane extracts isolated from whole TA muscles were layered on top of a linear 10–40% glycerol gradient, prepared in Buffer G (20 mM Tris, pH 7.6, 5 mM EDTA, pH 7.4, 100 mM KCl, 1 mM DTT, 0.25% sodium deoxycholate, and 1 mM sodium orthovanadate). Following centrifugation at 131,300 × g for 24 hr at 4 °C using a MLS-50 Swinging-Bucket rotor (Beckman Coulter, Brea, CA) to sediment protein complexes, 250 μl fractions were collected from the bottom of the tube, and alternate fractions were subjected to TCA precipitation (10% TCA was added to samples) for 24 hr at 4 °C. Precipitates were resuspended and analyzed by SDS-PAGE and immunoblotting.

### Immunoblotting and immunoprecipitation

For immunoprecipitation, muscle homogenates or glycerol gradient fractions were incubated with specific antibodies (control sample contained 1 μg of non-specific IgG) overnight at 4 °C on a rotating device, and then Protein A/G agarose was added for additional 4 hr at 4 °C. To remove nonspecific or weakly bound proteins, precipitates were washed with 10 bed volumes of the following buffers: high (50 mM Tris-HCl, pH 8, 500 mM NaCl, 0.1% SDS, 0.1% Triton, 5 mM EDTA), medium (50 mM Tris-HCl, pH 8, 150 mM NaCl, 0.1% SDS, 0.1% Triton, 5 mM EDTA), and low (50 mM Tris-HCl, pH 8, 0.1% Triton, 5 mM EDTA) salt buffers. Protein precipitates were then analyzed by SDS-PAGE and immunoblotting.

For immunoblotting, samples were resolved by SDS-PAGE, transferred onto PVDF membrane, and immunoblotted with specific primary antibodies, and sequentially with secondary antibodies conjugated to HRP (anti heavy and light chains specific antibodies, or anti light chain specific antibody).

### Immunofluorescence staining of frozen muscle sections

Cross-sections of mouse TA (20 μm) were fixed in 4% paraformaldehyde (PFA), and incubated in blocking solution (0.2% bovine serum albumin and 5% normal goat serum in PBS-T) for 1 hr at room temperature. Immunostaining was performed using primary antibodies (1:50), and secondary antibodies conjugated to Alexa 568 or Alexa 679 (1:1000). Images were collected with an inverted LSM 710 laser scanning confocal microscope (Zeiss, Jena, Germany) with a Plan-Apochromat ×60 1.4 NA objective lens and BP 593/46, BP 525/50, and 640–797 filters, and analyzed with Imaris 8.2 software. For Man2b1 staining, frozen sections from normal muscles (Fig S2a) or ones electroporated with GFP-LC3 plasmid (5 d after electroporation mice were starved for 1 or 2 d) were fixed in 4% PFA and stained as described above. Sections were imaged using the Elyra 7 eLS lattice SIM super resolution microscope by Zeiss with a pco.edge sCMOS camera. A X63 1.46 NA oil immersion objective with 561nm and 642nm lasers. 16-bit 2D image data sets were collected with 13 phases. The SIM^2 image processing tool by Zeiss was used.

For MYH7 staining, frozen muscle sections were fixed in 4% PFA. Then, antigen retrieval was performed by soaking sections in 10 mM citrate buffer (containing 1.8 mM Citric Acid and 8.2 mM Sodium Citrate), heating for 20 min in microwave (lowest heat level), and letting cool inside the buffer to room temperature. Following three washes with PBST, samples were incubated in blocking solution (10% goat serum in PBST) for 1 hr at room temperature, and then with MYH7 antibody (1:5000) overnight at 4°C, and a secondary antibody conjugated to Alexa 568 (1:400).

### *In vivo* [^3^H]-2-deoxyglucose uptake by the skeletal muscle

*In vivo* glucose uptake was performed as described before ^686^. Mice were fasted for 2 days and anesthetized with sodium pentobarbital. After 25 min, blood was collected from tail to determine basal glucose and background radioactivity. [^3^H]-2-deoxyglucose (0.33 μCi/gram BW) was then injected retro-orbitally, and blood was collected every 5-10 min up to 45 min for glucose and [^3^H]-2-deoxyglucose measurements. Mice were euthanized and contralateral TA muscles were carefully removed, weighted, and frozen in liquid N2. To assess glucose uptake, dissected TA muscles were homogenized in lysis buffer (20 mM Tris, pH 7.4, 5 mM EDTA, 10 mM sodium pyrophosphate, 100 mM NaF, 2 mM sodium orthovanadate, 10 μg/ml aprotinin, 10 μg/ml leupeptin, 3 mM benzamidine, and 1 mM PMSF), and accumulation of radioactivity (total and phosphorylated [^3^H]-2-deoxyglucose) in muscle was assessed in muscle homogenates using scintillation counter and was calculated as described ^68^.

### Quantitative real-time PCR

Total RNA was isolated from muscle using TRI-reagent (Sigma, #T9424) and served as a template for synthesis of cDNA by reverse transcription (Quanta script cDNA synthesis kit, #84005, #84002). Real-time qPCR was performed on mouse target genes using specific primers (Table S2) and PerfecTa SYBR Green FastMix (Quanta 84071) according to manufacturer’s protocol.

### Oleic acid uptake by c2c12 myotubes

c2c12 cells were plated to reach 80-90% confluence on the following day in growth media (DMEM containing 10% FBS, 1% Penicillin/streptomycin, 1% glutamate) at 37 ⁰C and 5% CO2. Media was then replaced to differentiation media (DMEM, 2% Horse serum, 1% Penicillin/streptomycin, 1% glutamate), which was refreshed every other day. On day 4, when myotubes were clearly visible, cells were treated for 4 hr with 100 μM oleic acid (O1008-1g, sigma) and 29 μM BSA ^69^. To confirm oleic acid uptake, cells were stained with Oil Red O (O0625, Merck) as follows: untreated and treated myotubes were washed with PBS, fixed with 4% PFA for 20 min, washed with 60% Isopropanol and stained with Oil Red O stain ^70^. Then, images were taken with Leica DMI8 Inverted Fluorescent Microscope (20X). To determine *HEXA* and *MAN2B1* gene expression in treated and untreated cells, total RNA was isolated by TRIzol Reagent (Ambion-Cat.No:15596018) and 2 ug RNA served as a template for synthesis of cDNA by reverse transcription (Agentek Cat.No:95047-100). Real-time qPCR was performed on mouse target genes using specific primers (Table S2) and SYBR Green Master Mix (Applied Biosystems - 4367659).

### Chromatin immunoprecipitation (ChIP) Assay

The ChIP assay was performed using the Millipore ChIP Assay Kit (Cat# 17-295). Four TA muscles per experimental condition (fed and fasted mice) were lysed in Nuclear Extraction Buffer (10 mM HEPES pH 7.4, 10 mM KCl, 5 mM MgCl2, 0.5 mM DTT, protease and phosphatase inhibitors) with the QIAGEN Tissue Lyser II (30 Hz, 48 sec) and pooled. After crosslinking with formaldehyde (1% v/v final concentration) for 10 min at 4°C, crosslinking was quenched with 1.25 mM glycine for 5 min at room temperature and the samples were centrifuged at 1000 x g, at 4°C for 5 min. Pellets were resuspended in SDS lysis buffer (Cat# 20-163) and sonicated with the Covaris E220 ultrasonicator (Peak Power 75, Duty Factor 26, Cycles/Burst 200, Temperature 6°C, 960 sec) to fragment DNA. The samples were then centrifuged at 1000 x g, 4°C for 5 min, and the supernatant was diluted with ChIP Dilution Buffer (Cat# 20-153) and precleared for 1 hr at 4°C with 5 μl Protein A Agarose/Salmon Sperm DNA (Cat# 16-157C), then incubated with Protein A Agarose/Salmon Sperm DNA and 5 μg human IgG (Sigma, Cat# I4506) or PPAR-γ Antibody (E-8) (Santa Cruz, Cat# sc-7273) overnight at 4°C. The beads were washed with the buffers supplied in the kit in the following order for 7 min each: Low Salt Immune Complex Wash Buffer (Cat# 20-154), High Salt Immune Complex Wash Buffer (Cat# 20-155), LiCl Immune Complex Wash Buffer (Cat# 20-156), TE Buffer (Cat# 20-157). DNA was eluted in Elution Buffer (1% SDS, 0.1 M NaHCO3) with 0.2 mg/ml Proteinase K for 5 hr with shaking at 1200 rpm, 65°C. Eluted DNA was purified by isopropanol precipitation and subjected to qPCR analysis. Fold enrichment was calculated by dividing Cq values from the PPAR-γ immunoprecipitation by IgG signal.

### Isolation of glycosylated proteins from membrane extracts

Membrane extracts isolated from whole TA muscles were incubated each with 100 μl Sambucus Nigra Lectin-bound beads (SNA, EBL)(50% resin slurry) for 30 min at room temperature on a rotating device. Following centrifugation for 5 min at 1,000 x g, protein precipitates were washed three times with wash buffer (20 mM Tris pH 7.2, 5 mM EGTA, 100 mM KCl, 1% Triton X-100, 10 mM Sodium Pyrophosphate, 50 mM NaF, 2 mM Sodium Orthovanadate, 10 μg/ml Aprotinin, 10 μg/ml Leupeptin, 3 mM Benzamidine, and 1 mM PMSF), and analyzed by SDS-PAGE and immunoblotting.

### *In vitro* deglycosylation assay

Equal amounts (25 μg) of whole normal muscle extracts were incubated with 1 x Glycoprotein denaturing buffer (Biolabs) at 100 °C for 10 min to denature glycoproteins, and the deglycosylation reaction was performed according to manufacturer instructions. Specifically, *in vitro* deglycosylation was performed in 40 μl mixtures containing the denatured muscle extracts, 1% NP-40, and PNGase F (500U)(Cat# P7367, Biolabs) in 1 x GlycoBuffer-2 (Biolabs). After incubation for 60 min at 37 °C, samples were analyzed by SDS-PAGE and immunoblotting.

### Statistical analysis and image acquisition

All data are presented as means ± SEM. The statistical significance was accessed with one-tailed paired Student’s *t* test when one limb was compared to the contralateral limb in the same fasted mouse, and one-tailed unpaired Student’s *t* test when two groups of mice were compared. When several groups of mice were compared ANOVA test was used. Muscle sections used for fiber size analysis were imaged at room temperature with a Nikon Ni-U upright fluorescence microscope with Plan Fluor 10× 0.3-NA or Plan Fluor 20x 0.5NA objective lenses and a Hamamatsu C8484-03 cooled CCD camera. For immunofluorescence experiments presented in Fig. 3b and f, images were collected with an inverted LSM 710 laser scanning confocal microscope (Zeiss, Jena, Germany) with a Plan-Apochromat ×60 1.4 NA objective lens. For the immunofluorescence experiment presented in Fig. 3a, images were collected with a Nikon Ni-U upright fluorescence microscope and a Hamamatsu C8484-03 cooled CCD camera. GFP was detected using the Argon laser (488nm) and emission was collected by PMT detector with 525/50 nm filter (green). Alexa 555 and Alexa 647 were excited using 543nm and 639 nm lasers and emissions were collected by GaAsp detectors within BP 593/46 nm (red) and BP 718/156nm (far red), respectively. Image acquisition was performed using 3.2 ZEN imaging software (Zeiss), and data processing was performed using Imaris (Bitplane) or Metamorph (Molecular Devices) software.

Colocalization (PCC) and percent colocalization in region of interest (ROI is the region of cooccurrence) were quantified between signal intensities of β-dystroglycan and NAGLU (Fig. 3b) using confocal images, and image analysis was performed with Imaris (Bitplane, ver 9.1.2) by ImarisColoc module. This pixel-based module provided a cut-off threshold for separating signal from background pixels and defined the overlap between any two-color channels (colocalized pixels) of the image. Because PCC values were close to 1, we conclude that these proteins colocalize in skeletal muscle. Black and white images were processed with Adobe Photoshop CS5, version 12.1×64 software.

## Data availability

The materials and associated protocols generated and presented in the current study are available from the corresponding author on request.

## ACKNOWLEDGEMENTS

This project was supported by grants from the Israel Science Foundation (grant no. 1068/19), the Niedersachsen-Deutsche (grant no. ZN3008), and Israel Ministry of Health (grant no. 3-16061) to S. Cohen. Additional funds were received from the Russell Berrie Nanotechnology Institute, Technion to S. Cohen, and the Zuckerman STEM Leadership Program to J.E.G.

We are extremely thankful to Dr. Vered Padler-Karavani for providing valuable insights and suggestions on the glycosylation experiments. We also thank Dr. Yara Eid-Mutlak for helping with the *in vitro* deglycosylation assay (Fig. 2a), and the LS&E Microscopy Center at Technion for their assistance with the confocal microscope.

## AUTHOR CONTRIBUTIONS

S.J. and S.Z. performed all experiments. A.V. performed experiments presented in Fig. 3. D.K. performed the experiment presented as Fig. 4g. J.E.G. performed the experiments presented in Figs. 4b and S1c-d. A.F. performed the fiber size analyses and densitometric measurements. A.A. helped with the ChIP assay (Fig. 4b). S.J., S.Z, A.V., J.E.G., S.C designed experiments and analyzed data. S.C. wrote the paper.

---

**Figure S1.**
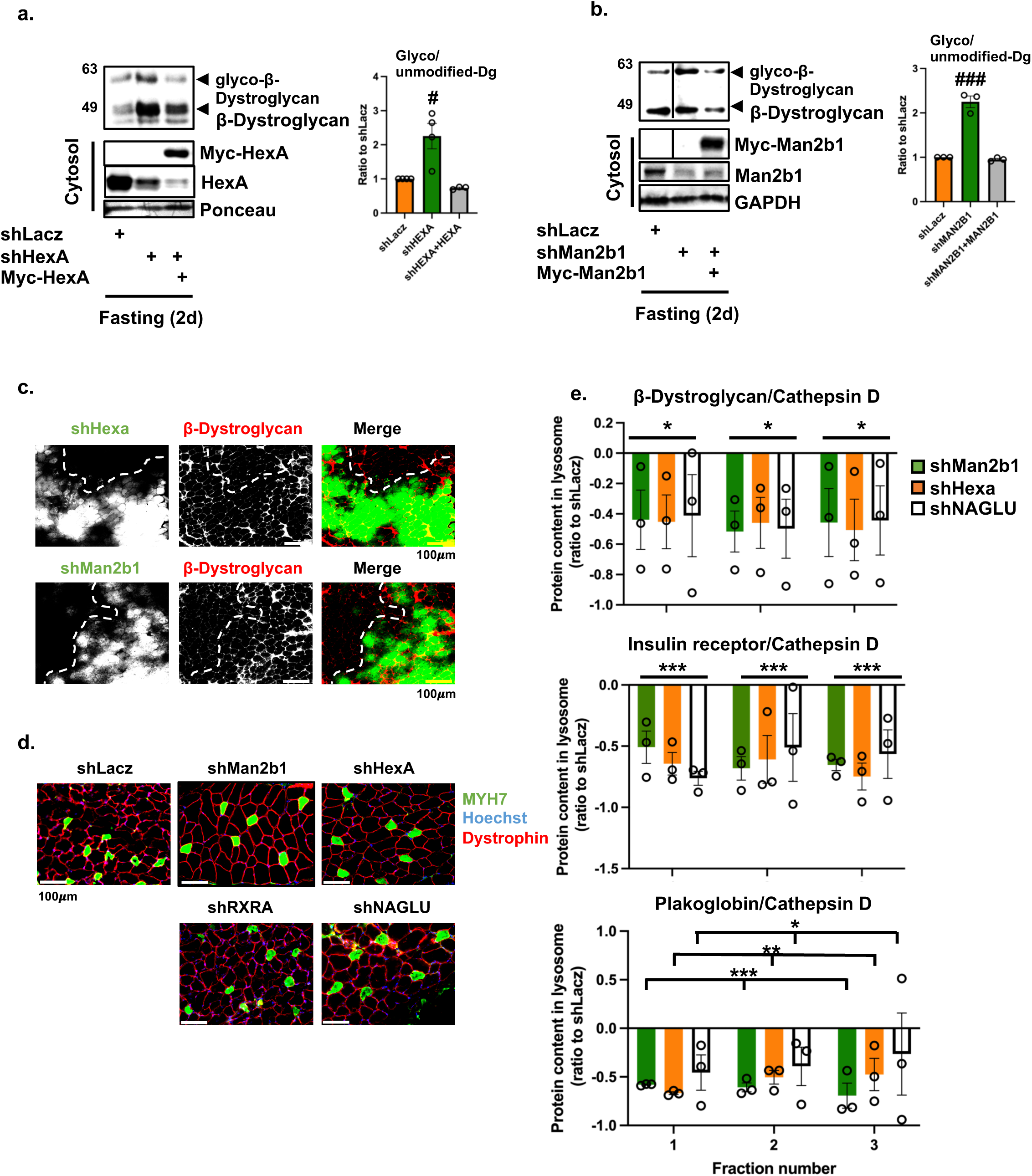
β-Dystroglycan accumulates on the membrane of atrophying muscles expressing shMan2b1 or shHexA. (a-b) During fasting, ectopic expression of Myc-Man2b1 or Myc-HexA reduces the accumulation of glycosylated β-dystroglycan imposed by shMan2b1 or shHexA. Equal amounts of membrane extracts from muscles expressing shHexA in the presence or absence of Myc-HexA (a), shMan2b1 in the presence or absence of Myc-Man2b1 (b) or shLacz from fasted mice were analyzed by immunoblotting. Graphs depict mean ratio to shLacz in fasting ± SEM. n=3-4 mice per condition. #, p<0.05; ###, p<0.0005 *vs.* fasting shLacz by ANOVA. The data are representative of three independent experiments. Black line indicates the removal of intervening lanes for presentation purposes. (c) During fasting, expression of shMan2b1 or shHexA markedly attenuates β-dystroglycan loss. Cross-sections of TA muscles expressing shMan2b1 (189/281 cells are transfected, green) or shHexA (207/282 cells are transfected, green) from fasted mice were stained with an antibody against β-dystroglycan (red). β-dystroglycan is lost in fibers, which do not express shMan2b1 or shHexA (average intensity of β-dystroglycan in non-transfected fibers is 11.996 and 13.297, respectively). Expression of shMan2b1 or shHexA markedly attenuated the loss of β-dystroglycan (average intensity of β-dystroglycan in transfected fibers is 41.458 and 33.158, respectively). Bar, 100 μm. The data are representative of two independent experiments. (d) Cross-sections of TA muscles expressing shLacz, shMan2b, shHexA, shRXRA or shNAGLU from fasted mice were stained with MYH7 (green) or dystrophin (red, for membrane staining) antibodies. Bar, 100 μm. The data are representative of two independent experiments. (e) Downregulation of Man2b1 or HexA reduces trafficking of β-dystroglycan, insulin receptor and plakoglobin to lysosomes. Densitometric measurements of the three heaviest Nycodenz fractions in blots presented in Fig. 2h. Ratios of each protein to the lysosomal marker cathepsin D are shown, and are plotted as the mean reduction in protein content in lysosome relative to shLacz control ± SEM. n=3 mice per condition. *, p<0.05; **, p<0.005, ***, p<0.0005 *vs.* shLacz by two-way ANOVA. Data are representative of three independent experiments.

**Figure S2.**
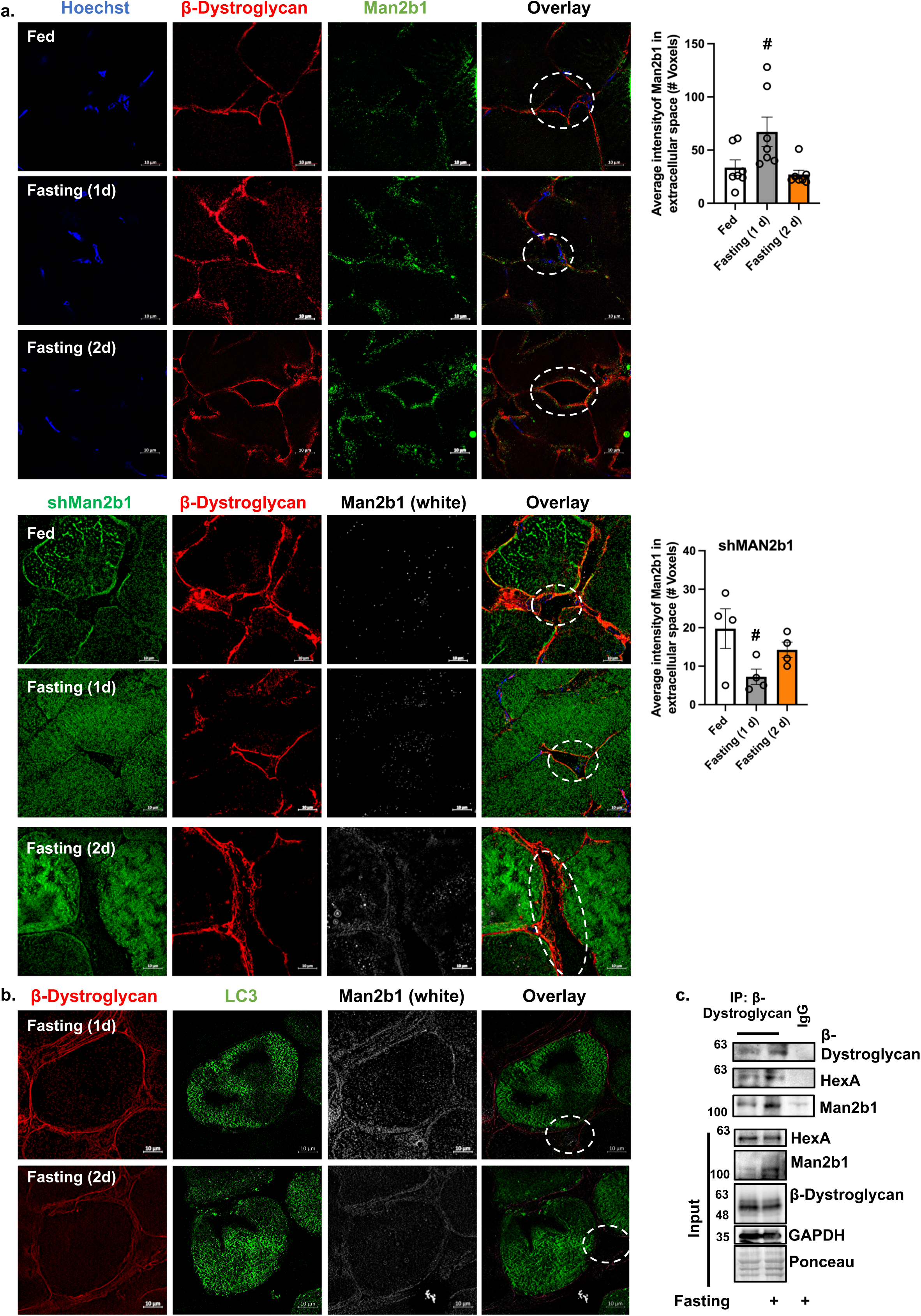
Man2b1 is secreted to the extracellular milieu at 1 d of fasting. Cross-sections of normal TA muscles from fed or fasted mice (1 or 2 d) (a), expressing shNAGLU (a) or GFP-LC3 (b) were stained with the indicated antibodies and imaged with super-resolution microscope (SIM). Extracellular areas are marked with a broken line. Bar, 10 μm. The data are representative of three independent experiments. Graphs depict mean intensity of Man2b1 protein in the extracellular space ± SEM. n=3 mice per condition. #, p<0.05 *vs.* fed by ANOVA. (c) Membrane extracts of muscles from fed or fasted mice were subjected to β-dystroglycan immunoprecipitation, and protein precipitates were analyzed by SDS-PAGE and immunoblotting. The data are representative of two independent experiments. n= 3 mice.

**Figure S3.**
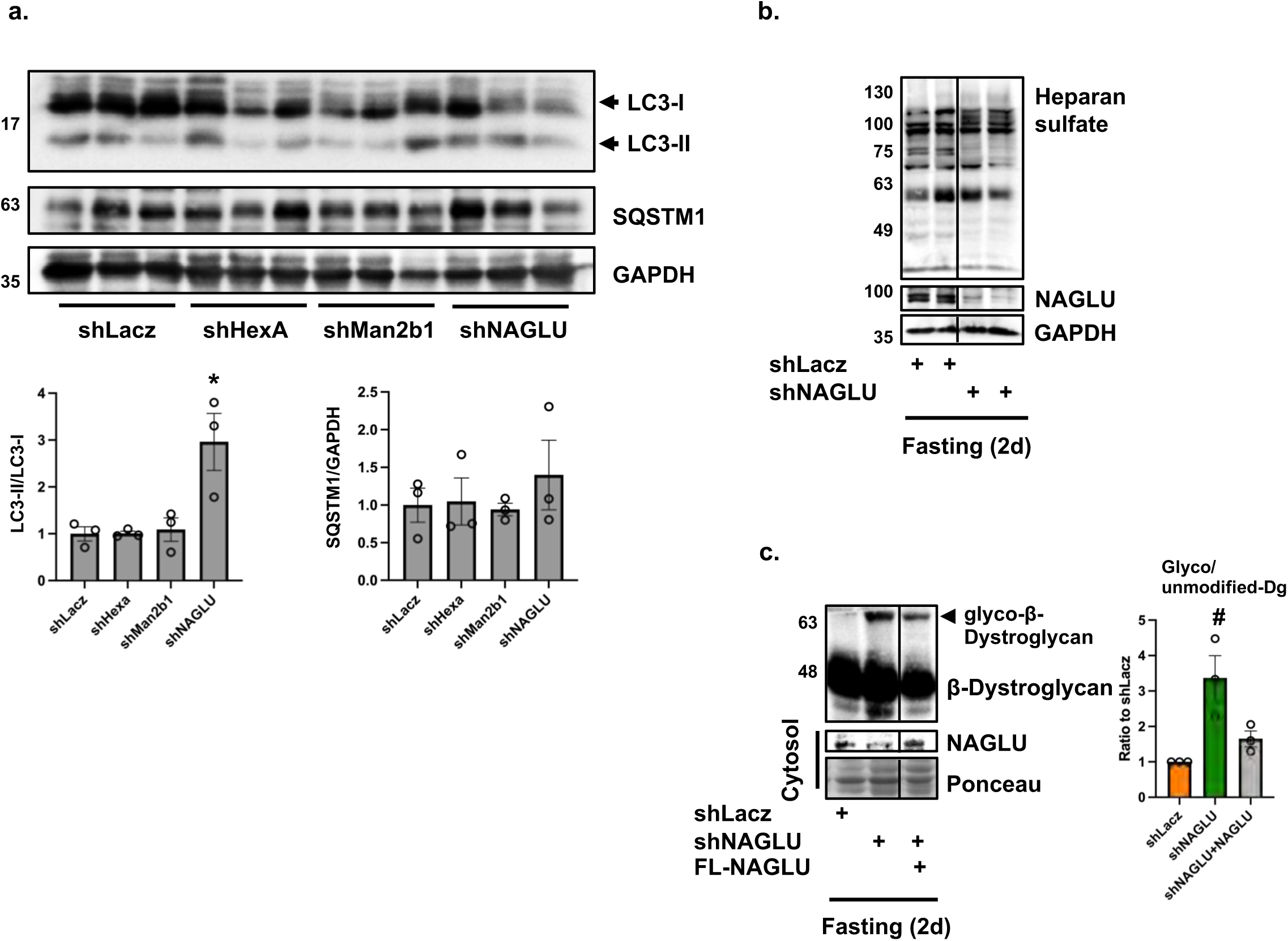
Downregulation of NAGLU in atrophying mouse muscle does not impair autophagy or lysosome function. (a) Homogenates of muscles expressing shLacz, shMan2b1, shHexA or shNAGLU from fasted mice (2 d) were analyzed by SDS-PAGE and immunoblot. Bottom: densitometric measurements of presented blots. Mean ratios of LC3-II/LC3-I and SQSTM1/GAPDH are presented. Data are plotted as the mean fold change relative to shLacz ± SEM. n=3 mice per condition. *, p<0.05 *vs.* shLacz by ANOVA. (b) Homogenates of muscles expressing shLacz or shNAGLU from fasted mice were analyzed by immunoblotting using heparan sulfate, NAGLU or GAPDH antibodies. Black lines indicate the removal of intervening lanes for presentation purposes. (c) During fasting, ectopic expression of full length NAGLU (FL-NAGLU) reduces the accumulation of glycosylated β-dystroglycan imposed by shNAGLU. Equal amounts of membrane extracts from muscles expressing shNAGLU in the presence or absence of FL-NAGLU or shLacz from fasted mice were analyzed by immunoblotting. Graph depicts mean ratio to shLacz ± SEM. n=3 mice per condition. #, p<0.05 *vs.* shLacz by ANOVA. The data are representative of three independent experiments. Black lines indicate the removal of intervening lanes for presentation purposes.

**Figure S4.**
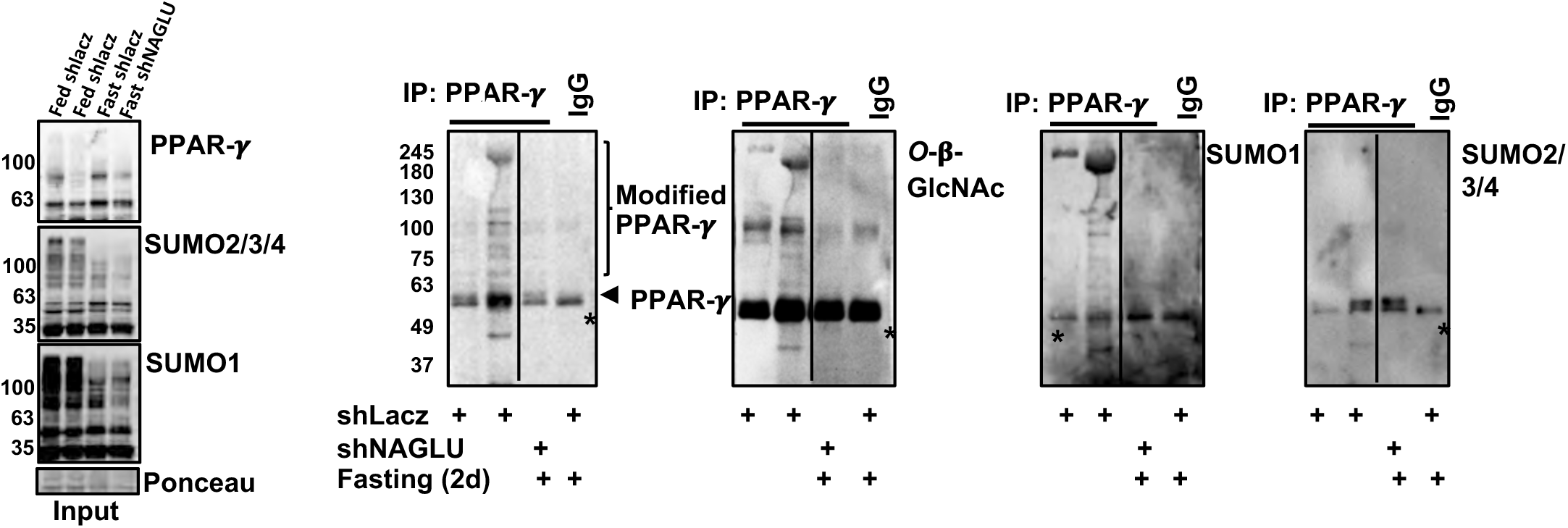
During fasting, NAGLU downregulation reduces *O*-β-GlcNAcylation of SUMOylated PPAR-γ. PPAR-γ *O*-β-GlcNAcylation in fasting requires NAGLU. PPAR-γ was immunoprecipitated from equal amounts of whole cell extracts from muscles expressing shNAGLU or shLacz from fed or fasted mice. Protein precipitates were analyzed by immunoblotting using antibodies against PPAR-γ, *O*-β-GlcNAcylation, SUMO1 or SUMO2/3/4. Asterisks represent nonspecific bands. The data are representative of two independent experiments. Black lines indicate the removal of intervening lanes for presentation purposes.

**Figure S5.**
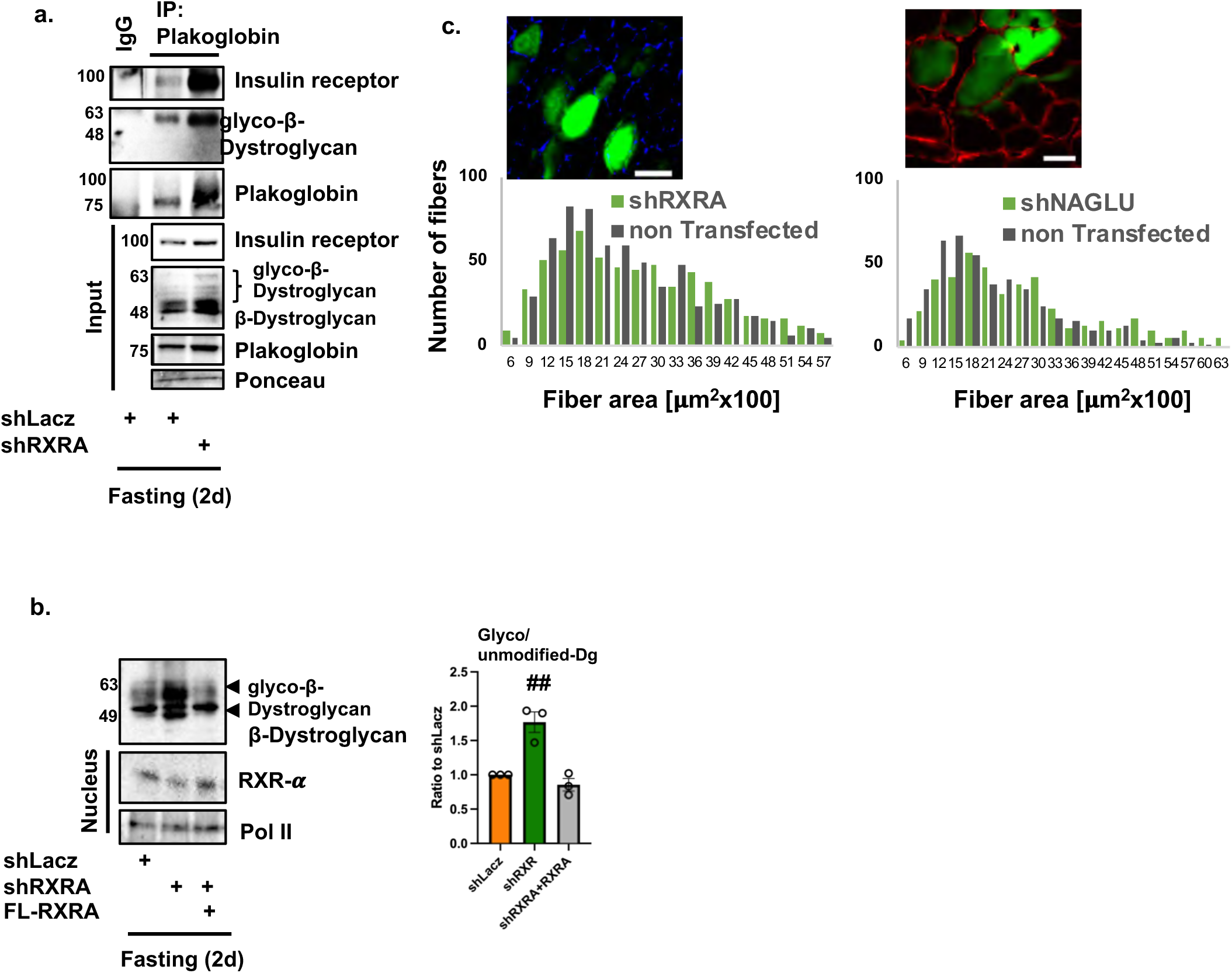
(a) Downregulation of RXR-α promotes accumulation of glycosylated β-dystroglycan-insulin receptor-plakoglobin assemblies on the muscle membrane. Plakoglobin immunoprecipitation from membrane extracts of whole TA muscles expressing shRXRA or shLacz from fasted mice. Protein precipitates and input samples were analyzed by immunoblotting. Ponceau S staining is shown as a loading control for input blots. The data are representative of two independent experiments. (b) During fasting, ectopic expression of full length RXR-α (FL-RXRA) reduces the accumulation of glycosylated β-dystroglycan imposed by shRXRA. Equal amounts of membrane extracts from muscles expressing shRXRA or shLacz from fasted mice were analyzed by immunoblotting. Graph depicts mean ratio to shLacz ± SEM. n=3 mice per condition. ##, p<0.005 *vs.* shLacz by ANOVA. The data are representative of two independent experiments. (c) Downregulation of RXR-α or NAGLU attenuates fiber atrophy during fasting. Measurements of cross-sectional areas of 670 fibers expressing shRXRA or 454 fibers expressing shNAGLU (green bars) *vs.* the same number of adjacent non-transfected fibers (dark bars). n=4 mice. Dystrophin staining is in red. Bar, 50μm.

**Table S1.**
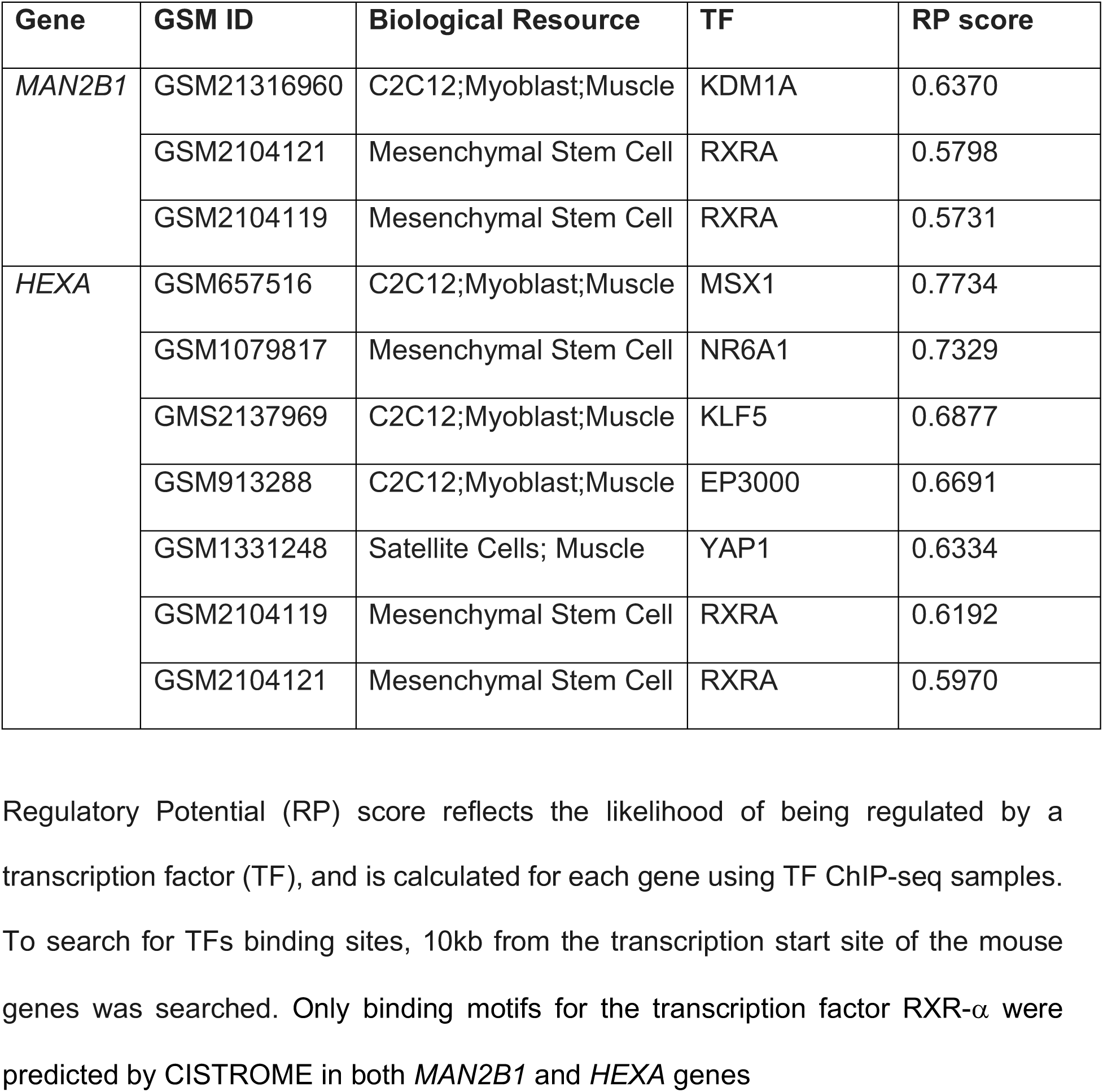
Potential transcription factor-binding sites in the promoter regions of *HEXA* and *MAN2B1* genes using the CISTROME portal.

**Table S2.**
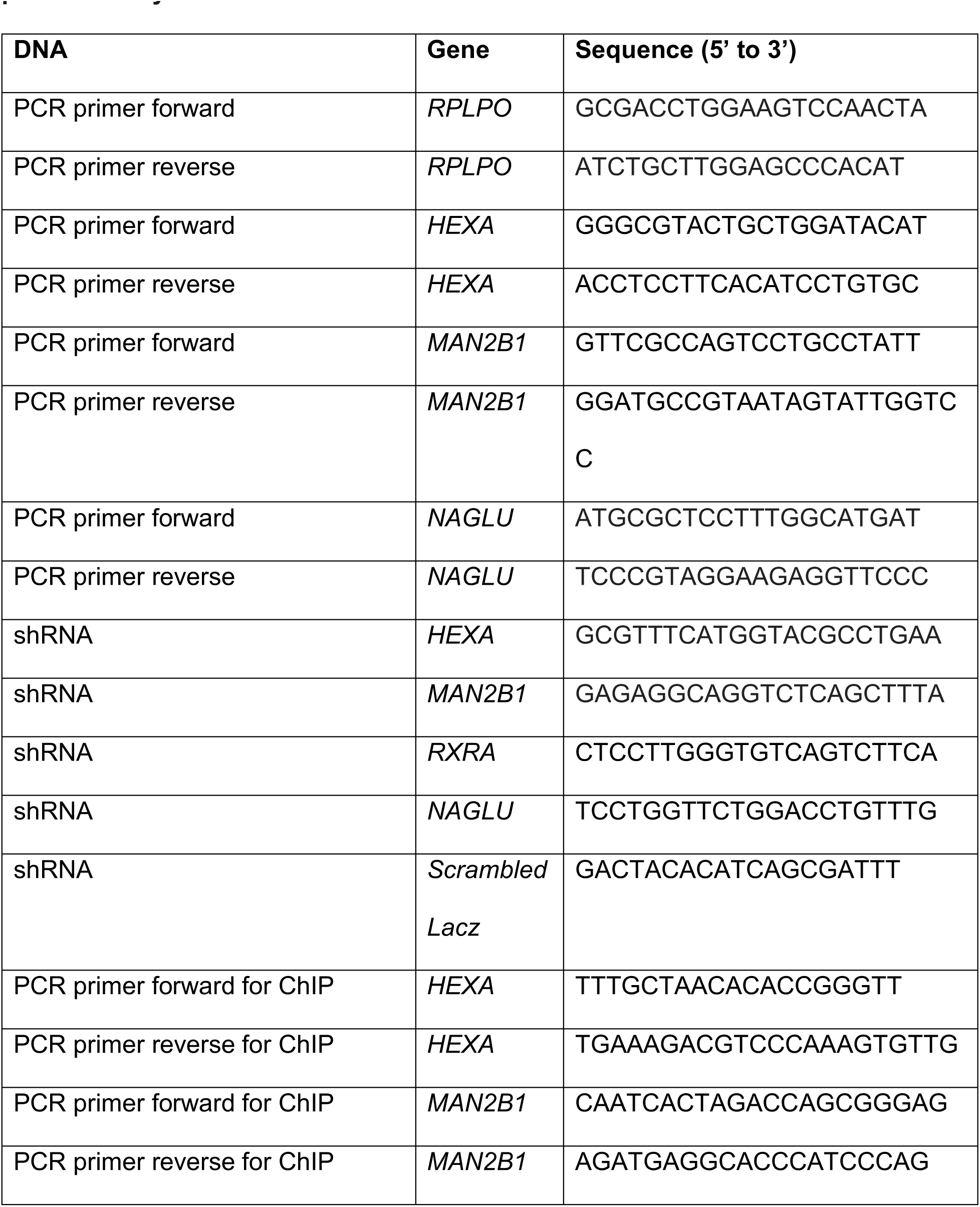
Quantitative PCR primers and shRNA oligonucleotides used in the present study.

## REFERENCES

1. Lee, J. & Pilch, P. F. The insulin receptor: Structure, function, and signaling. American Journal of Physiology - Cell Physiology Preprint at 10.1152/ajpcell.1994.266.2.c319 (1994).

2. Schiaffino, S. & Mammucari, C. Regulation of skeletal muscle growth by the IGF1-Akt/PKB pathway: Insights from genetic models. Skeletal Muscle Preprint at 10.1186/2044-5040-1-4 (2011).

3. Glass, D. J. Skeletal muscle hypertrophy and atrophy signaling pathways. Int J Biochem Cell Biol 37, 1974–1984 (2005).

4. Sandri, M. et al. Foxo transcription factors induce the atrophy-related ubiquitin ligase atrogin-1 and cause skeletal muscle atrophy. Cell 117, 399–412 (2004).

5. Latres, E. et al. Insulin-like growth factor-1 (IGF-1) inversely regulates atrophy-induced genes via the phosphatidylinositol 3-kinase/Akt/mammalian target of rapamycin (PI3K/Akt/mTOR) pathway. Journal of Biological Chemistry (2005) doi:10.1074/jbc.M407517200.

6. Eid Mutlak, Y., et al. A signaling hub of insulin receptor, dystrophin glycoprotein complex and plakoglobin regulates muscle size. Nat Commun 11, 1381 (2020).

7 De Meyts, P. The Insulin Receptor and Its Signal Transduction Network. Endotext (2000).

8. Birgisdottir, Å. B. et al. The LIR motif - crucial for selective autophagy. J Cell Sci (2013) doi:jcs.126128 [pii]\r10.1242/jcs.126128.

9. Grady, R. M. et al. Skeletal and cardiac myopathies in mice lacking utrophin and dystrophin: a model for Duchenne muscular dystrophy. Cell 90, 729–38 (1997).

10. Lapidos, K. A., Kakkar, R. & McNally, E. M. The Dystrophin Glycoprotein Complex. Circ Res 94, 1023–1031 (2004).

11. Hoffman, E. P., Brown Jr., R. H. & Kunkel, L. M. Dystrophin: the protein product of the Duchenne muscular dystrophy locus. Cell 51, 919–928 (1987).

12. Hoffman, E. P. & Kunkel, L. M. Dystrophin abnormalities in Duchenne/Becker muscular dystrophy. Neuron 2, 1019–1029 (1989).

13. Lauretani, F. et al. Age-associated changes in skeletal muscles and their effect on mobility: an operational diagnosis of sarcopenia. J Appl Physiol 95, 1851– 1860 (2003).

14. Ervasti, J. M. & Campbell, K. P. Membrane organization of the dystrophin-glycoprotein complex. Cell 66, 1121–1131 (1991).

15. Constantin, B. Dystrophin complex functions as a scaffold for signalling proteins. Biochim Biophys Acta 1838, 635–642 (2014).

16. Townsend, D. Finding the sweet spot: assembly and glycosylation of the dystrophin-associated glycoprotein complex. Anat Rec (Hoboken) 297, 1694– 705 (2014).

17. Hohenester, E. Laminin G-like domains: dystroglycan-specific lectins. Curr Opin Struct Biol 56, 56–63 (2019).

18. Manya, H. & Endo, T. Glycosylation with ribitol-phosphate in mammals: New insights into the O-mannosyl glycan. Biochim Biophys Acta Gen Subj 1861, 2462–2472 (2017).

19. Yoshida-Moriguchi, T. & Campbell, K. P. Matriglycan: a novel polysaccharide that links dystroglycan to the basement membrane. Glycobiology 25, 702–713 (2015).

20. Wells, L. The o-mannosylation pathway: glycosyltransferases and proteins implicated in congenital muscular dystrophy. J Biol Chem 288, 6930–6935 (2013).

21. Sheikh, M. O., Halmo, S. M. & Wells, L. Recent advancements in understanding mammalian O-mannosylation. Glycobiology 27, 806–819 (2017).

22. Oppizzi, M. L., Akhavan, A., Singh, M., Fata, J. E. & Muschler, J. L. Nuclear Translocation of β-Dystroglycan Reveals a Distinctive Trafficking Pattern of Autoproteolyzed Mucins. Traffic 9, 2063 (2008).

23. Barresi, R. & Campbell, K. P. Dystroglycan: From biosynthesis to pathogenesis of human disease. J Cell Sci (2006) doi:10.1242/jcs.02814.

24. Olsson, G. M., Svensson, I., Zdolsek, J. M. & Brunk, U. T. Lysosomal enzyme leakage during the hypoxanthine/xanthine oxidase reaction. Virchows Arch B Cell Pathol Incl Mol Pathol (1988) doi:10.1007/BF02890041.

25. Birnie, G. D. Iodinated density gradient media: a practical approach: Edited by D. Rickwood IRL Press; Oxford and Washington, DC, 1983 xii + 240 pages. £8.50, $17.00. FEBS Lett (1984) doi:10.1016/0014-5793(84)80236-7.

26. Pertoft, H., Warmegard, B. & Hook, M. Heterogeneity of lysosomes originating from rat liver parenchymal cells. Metabolic relationship of subpopulations separated by density gradient centrifugation. Biochemical Journal (1978) doi:10.1042/bj1740309.

27. Mammucari, C. et al. FoxO3 controls autophagy in skeletal muscle in vivo. Cell Metab 6, 458–471 (2007).

28. Zhao, J. et al. FoxO3 coordinately activates protein degradation by the autophagic/lysosomal and proteasomal pathways in atrophying muscle cells. Cell Metab 6, 472–483 (2007).

29. Rickwood, D., Ford, T. & Graham, J. Nycodenz: A new nonionic iodinated gradient medium. Anal Biochem 123, 23–31 (1982).

30. Sawant, P. L., Shibko, S., Kumta, U. S. & Tappel, A. L. Isolation of rat-liver lysosomes and their general properties. Biochimica et Biophysica Acta (BBA) - Specialized Section on Enzymological Subjects 85, 82–92 (1964).

31. Elbaz, Y. & Schuldiner, M. Staying in touch: the molecular era of organelle contact sites. Trends Biochem Sci 36, 616–623 (2011).

32. Todkar, K., Ilamathi, H. S. & Germain, M. Mitochondria and Lysosomes: Discovering Bonds. Front Cell Dev Biol 5, 106 (2017).

33. Mizushima, N. Autophagy: process and function. Genes Dev 21, 2861–73 (2007).

34. Milan, G. et al. Regulation of autophagy and the ubiquitin–proteasome system by the FoxO transcriptional network during muscle atrophy. Nat Commun 6, 6670 (2015).

35. Ju, J. S., Varadhachary, A. S., Miller, S. E. & Weihl, C. C. Quantitation of ‘autophagic flux’ in mature skeletal muscle. Autophagy (2010) doi:10.4161/auto.6.7.12785.

36. Grewal, P. K. & Hewitt, J. E. Glycosylation defects: a new mechanism for muscular dystrophy? Hum Mol Genet (2003) doi:10.1093/hmg/ddg272.

37. Wellen, K. E. & Thompson, C. B. Cellular Metabolic Stress: Considering How Cells Respond to Nutrient Excess. Molecular Cell Preprint at 10.1016/j.molcel.2010.10.004 (2010).

38. Je, G., et al. A semiautomated measurement of muscle fiber size using the Imaris software. Am J Physiol Cell Physiol 321, C615–C631 (2021).

39. Kleijmeer, M. et al. Reorganization of multivesicular bodies regulates MHC class II 11 antigen presentation by dendritic cells. Journal of Cell Biology (2001) doi:10.1083/jcb.200103071.

40. Schmidtchen, A. et al. NAGLU mutations underlying Sanfilippo syndrome type B. Am J Hum Genet (1998) doi:10.1086/301685.

41. Lotfi, P. et al. Trehalose reduces retinal degeneration, neuroinflammation and storage burden caused by a lysosomal hydrolase deficiency. Autophagy (2018) doi:10.1080/15548627.2018.1474313.

42. Liu, T. et al. Cistrome: An integrative platform for transcriptional regulation studies. Genome Biol (2011) doi:10.1186/gb-2011-12-8-r83.

43. Kliewer, S. A., Umesono, K., Noonan, D. J., Heyman, R. A. & Evans, R. M. Convergence of 9-cis retinoic acid and peroxisome proliferator signalling pathways through heterodimer formation of their receptors. Nature (1992) doi:10.1038/358771a0.

44. Kintscher, U. & Law, R. E. PPARγ-mediated insulin sensitization: The importance of fat versus muscle. American Journal of Physiology - Endocrinology and Metabolism Preprint at 10.1152/ajpendo.00440.2004 (2005).

45. Wolins, N. E. et al. OXPAT/PAT-1 Is a PPAR-Induced Lipid Droplet Protein That Promotes Fatty Acid Utilization. Diabetes 55, 3418–3428 (2006).

46. Phua, W. W. T., Wong, M. X. Y., Liao, Z. & Tan, N. S. An aPPARent Functional Consequence in Skeletal Muscle Physiology via Peroxisome Proliferator-Activated Receptors. International Journal of Molecular Sciences 2018, Vol. 19, Page 1425 19, 1425 (2018).

47. Høstmark, A. T. & Haug, A. Percentages of oleic acid and arachidonic acid are inversely related in phospholipids of human sera. Lipids Health Dis 12, 106 (2013).

48. Ji, S., Park, S. Y., Roth, J., Kim, H. S. & Cho, J. W. O-GlcNAc modification of PPARγ reduces its transcriptional activity. Biochem Biophys Res Commun (2012) doi:10.1016/j.bbrc.2011.12.086.

49. Yang, X. & Qian, K. Protein O-GlcNAcylation: Emerging mechanisms and functions. Nature Reviews Molecular Cell Biology Preprint at 10.1038/nrm.2017.22 (2017).

50. Brunmeir, R. & Xu, F. Functional Regulation of PPARs through Post-Translational Modifications. Int J Mol Sci 19, (2018).

51. Mikkonen, L., Hirvonen, J. & Jänne, O. A. SUMO-1 regulates body weight and adipogenesis via PPARγ in male and female mice. Endocrinology 154, 698– 708 (2013).

52. Winchester, B. Lysosomal metabolism of glycoproteins. Glycobiology Preprint at 10.1093/glycob/cwi041 (2005).

53. Klaver, E. et al. Selective inhibition of N-linked glycosylation impairs receptor tyrosine kinase processing. DMM Disease Models and Mechanisms (2019) doi:10.1242/dmm.039602.

54. Heidenreich, K. A. & Brandenburg, D. Oligosaccharide heterogeneity of insulin receptors. comparison of N-linked glycosylation of insulin receptors in adipocytes and brain. Endocrinology (1986) doi:10.1210/endo-118-5-1835.

55. Collier, E. & Gorden, P. O-linked oligosaccharides on insulin receptor. Diabetes (1991) doi:10.2337/diabetes.40.2.197.

56. Stewart, M. D. & Sanderson, R. D. Heparan sulfate in the nucleus and its control of cellular functions. Matrix Biology Preprint at 10.1016/j.matbio.2013.10.009 (2014).

57. Nugent, M. A., Zaia, J. & Spencer, J. L. Heparan sulfate-protein binding specificity. Biochemistry (Moscow) Preprint at 10.1134/S0006297913070055 (2013).

58. Hardivillé, S. & Hart, G. W. Nutrient regulation of signaling, transcription, and cell physiology by O-GlcNAcylation. Cell Metabolism Preprint at 10.1016/j.cmet.2014.07.014 (2014).

59. Tawo, R. et al. The Ubiquitin Ligase CHIP Integrates Proteostasis and Aging by Regulation of Insulin Receptor Turnover. Cell (2017) doi:10.1016/j.cell.2017.04.003.

60. van der Crabben, S. N. et al. Prolonged fasting induces peripheral insulin resistance, which is not ameliorated by high-dose salicylate. Journal of Clinical Endocrinology and Metabolism (2008) doi:10.1210/jc.2006-2491.

61. Soll, A. H., Kahn, C. R. & Neville, D. M. Insulin binding to liver plasm membranes in the obese hyperglycemic (ob/ob) mouse. Demonstration of a decreased number of functionally normal receptors. J Biol Chem 250, 4702– 4707 (1975).

62. Haruta, T. et al. Amplification and analysis of promoter region of insulin receptor gene in a patient with leprechaunism associated with severe insulin resistance. Metabolism 44, 430–437 (1995).

63. Si, T., et al. Mutations in the insulin receptor gene in patients with genetic syndromes of insulin resistance. Adv Exp Med Biol 293, 721–730 (1991).

64. Choi, E., Zhang, X., Xing, C. & Yu, H. Mitotic Checkpoint Regulators Control Insulin Signaling and Metabolic Homeostasis. Cell 166, 567–581 (2016).

65. Goldbraikh, D. et al. USP1 deubiquitinates Akt to inhibit PI3K-Akt-FoxO signaling in muscle during prolonged starvation. EMBO Rep. (2020) doi:10.15252/embr.201948791.

66. Aweida, D., Rudesky, I., Volodin, A., Shimko, E. & Cohen, S. GSK3-β promotes calpain-1-mediated desmin filament depolymerization and myofibril loss in atrophy. J Cell Biol 217, 3698–3714 (2018).

67. Cohen, S. et al. During muscle atrophy, thick, but not thin, filament components are degraded by MuRF1-dependent ubiquitylation. J Cell Biol 185, 1083–1095 (2009).

68. Hinkley, J. M. et al. Constitutively Active CaMKKalpha Stimulates Skeletal Muscle Glucose Uptake in Insulin Resistant Mice In Vivo. Diabetes (2013) doi:10.2337/db13-0452.

69. Sanderson, L. M. et al. Peroxisome Proliferator-Activated Receptor β/δ (PPARβ/δ) but Not PPARα Serves as a Plasma Free Fatty Acid Sensor in Liver. Mol Cell Biol 29, 6257 (2009).

70. Liu, Y. et al. miR-324-5p Inhibits C2C12 cell Differentiation and Promotes Intramuscular Lipid Deposition through lncDUM and PM20D1. Mol Ther Nucleic Acids 22, 722–732 (2020).

